# Integrating multiple imputation with harmonization in multisite neuroimaging data: implications for site effects and biological inference

**DOI:** 10.64898/2026.11.14.766661

**Authors:** Yuxin Ren, Yiwen Dong, Davneet Minhas, Ann D. Cohen, Qiong Wu, Dana L. Tudorascu, The Alzheimer’s Disease Neuroimaging Initiative

## Abstract

Multisite neuroimaging harmonization in Alzheimer’s disease research with missing data has received limited attention in the literature. Here, we evaluated the performance of several harmonization methods in conjunction with multiple imputation (MI) to address missing data, using baseline [^18^F]flortaucipir tau positron emission tomography (PET) data from the Alzheimer’s Disease Neuroimaging Initiative (ADNI). We compared complete case and MICE-imputed datasets (*m* = 5) across the original ADNI-preprocessed data and four statistical harmonization approaches: ComBat with and without covariates, quantile normalization, and quantile regression. Weighted average standardized uptake value ratios (SUVRs) across six Braak regions were analyzed for each dataset-method combination. We assessed harmonization effectiveness using covariate-adjusted intraclass correlation coefficients and site-level pairwise differences, and evaluated biological effect preservation using Hedges’ *g*, regression coefficients, and statistical significance for comparisons between cognitively impaired and unimpaired participants. For imputed analyses, effect sizes and regression coefficients were pooled using Rubin’s rules, with statistical inference based on the pooled estimates. Post-processing harmonization reduced site-level variability compared to the original data, with method performance varying across evaluation metrics, while overall harmonization quality remained comparable after MI. Disease group differences remained statistically significant across all combinations, exhibiting a consistent regional pattern, strongest in Braak I and weakest in Braak VI. MI modestly increased effect size and coefficient estimates, with improved precision reflected by median standard error ratios below 1 (0.89 for effect sizes and 0.93 for coefficients). These findings indicate that integrating MI with harmonization preserves disease group differences and improves statistical efficiency without compromising harmonization performance, supporting a practical framework for multisite neuroimaging studies.

## 1 Introduction

Multisite neuroimaging studies provide important opportunities to investigate neurological and psychiatric disorders in large and heterogeneous populations. By pooling data collected across institutions, these studies can enhance generalizability through greater population diversity, increase statistical power through larger sample sizes, and ultimately improve the reproducibility of scientific findings. Large collaborative efforts, including the Alzheimer’s Disease Neuroimaging Initiative (ADNI) (Mueller et al., 2005), the Adolescent Brain Cognitive Development (ABCD) Study, the Human Connectome Project (HCP), and the Enhancing Neuro Imaging Genetics through Meta-Analysis (ENIGMA) Consortium, have illustrated the scientific value of this paradigm. However, combining neuroimaging data acquired across sites presents important methodological challenges, as differences in scanners, acquisition protocols, image reconstruction, and preprocessing can introduce systematic variation unrelated to the biological processes of interest (Jagust et al., 2024). These site-related differences, often referred to as batch effects, if not adequately addressed, can obscure biological associations, bias parameter estimates, and reduce the reliability and statistical efficiency of downstream analyses (Harrison et al., 2024). Such batch effects have been reported across imaging modalities. In structural magnetic resonance imaging (MRI), they have been shown to affect cortical thickness measurements (Han et al., 2006) and reduce statistical power in case-control analyses (Radua et al., 2020); similar site-related biases have been observed in diffusion MRI (Fortin et al., 2017), functional MRI (Yu et al., 2018), and positron emission tomography (PET) (Joshi et al., 2009). Across pooled neuroimaging datasets, these effects can be strong enough that individual scans can be assigned to their study of origin with high accuracy (Wachinger et al., 2021). Site effects are of particular concern for tau PET, as the U.S. Food and Drug Administration (FDA) approval of [^18^F]flortaucipir for clinical use (Fleisher et al., 2020) is expected to expand its acquisition across diverse clinical sites and scanners.

To address these challenges, statistical harmonization methods aim to mitigate technical variation while preserving biologically meaningful information across individuals and sites (Bayer, Thompson, et al., 2022). Harmonization can be performed at different stages of the neuroimaging pipeline, ranging from standardization of image acquisition and preprocessing (Landau et al., 2025) to statistical adjustment of derived imaging features. At the feature level, ComBat (Johnson et al., 2007) is one of the most widely used approaches for statistical harmonization. It models site effects as additive and multiplicative shifts in mean and variance, using empirical Bayes estimation to stabilize site-specific parameters. ComBat has been successfully applied across a range of neuroimaging modalities (Eshaghzadeh Torbati et al., 2021; Fortin et al., 2017, 2018; Radua et al., 2020; Yang et al., 2024; Yu et al., 2018), with extensions to nonlinear covariate modeling and Bayesian formulations (Pomponio et al., 2020; Reynolds et al., 2023). Alternative distribution-based approaches, such as quantile normalization (QN) and quantile regression (QR), provide greater flexibility to address distributional differences across sites (Zhang et al., 2026). While QN aligns empirical quantiles to enforce identical marginal distributions (Bolstad et al., 2003), QR can incorporate covariates when modeling site-specific differences at selected quantiles of the distribution (Cade & Noon, 2003; R. Koenker & Hallock, 2001). These approaches differ in the aspects of site-related variation they target and, consequently, may differ in their ability to reduce technical heterogeneity while preserving biologically meaningful variation.

An additional challenge that has received considerably less attention in multisite neuroimaging harmonization is incomplete covariate information. Most existing harmonization methods are developed for settings in which the imaging measures and biological covariates included in the harmonization model are completely observed. In practice, however, demographic, genetic, and clinical variables are frequently missing in large observational neuroimaging studies because of incomplete assessments, unavailable measurements, or participant attrition (Campos et al., 2015). Restricting analyses to participants with complete covariate information can substantially reduce the available sample size and may introduce bias when missingness is related to observed participant characteristics (Rubin, 1976), a scenario commonly encountered in large neuroimaging consortia. Multiple imputation (MI) provides a principled alternative by generating multiple plausible values for missing observations based on the observed data and appropriately accounting for the resulting uncertainty in statistical inference (Rubin, 1987; White et al., 2011). Although MI has been increasingly used to address missing clinical and demographic information in neuroimaging studies (Aracri et al., 2025; Chandrasekaran et al., 2024), its integration with multisite harmonization remains relatively understudied. In particular, it remains unclear whether harmonization methods retain their ability to reduce site-related variability when applied after multiple imputation and whether biologically meaningful associations are preserved in subsequent analyses.

In this study, we evaluated integrating multiple imputation with harmonization for multisite tau PET imaging with incomplete covariate data. Using baseline [^18^F]flortaucipir tau PET data from ADNI, we applied multivariate imputation by chained equations (MICE) to impute missing demographic, genetic, and clinical covariates. MICE imputes missing values through a series of conditional models and accommodates mixed data types (van Buuren & Groothuis-Oudshoorn, 2011; White et al., 2011). We compared complete-case (CC) and multiply imputed analyses across four harmonization approaches: ComBat with and without covariate adjustment, quantile normalization, and quantile regression. We evaluated the performance of each analysis strategy along two complementary dimensions: the reduction of site-related variability and the preservation of biologically meaningful differences between cognitively impaired and unimpaired participants across Braak regions. Through these comparisons, we aimed to determine whether multiple imputation can be effectively integrated with existing harmonization approaches while maintaining harmonization performance and downstream biological inference.

Our work contributes to the existing literature by providing a specific framework for addressing missing data and technical variability in multisite neuroimaging studies, together with a comprehensive evaluation of harmonization methods on data with missing values before and after MI. Furthermore, combining multiple imputation via MICE with data harmonization for tau PET imaging represents a novel contribution to multisite Alzheimer’s neuroimaging studies.

## 2 Methods

### 2.1 Data source

Baseline [^18^F]flortaucipir (FTP) tau PET data were obtained from the ADNI database (https://adni.loni.usc.edu/) on April 09, 2025. PET images were preprocessed using the standardized ADNI pipeline. Dynamic frames were first realigned for motion correction and averaged to create a single static image. The images were then standardized in orientation and voxel size and smoothed using scanner-specific Gaussian filters to achieve a common effective spatial resolution of 6 mm, reducing resolution-related differences across scanners (Joshi et al., 2009). Regional SUVRs were subsequently derived using each participant’s structural MRI and FreeSurfer-defined regions, with the inferior cerebellum used as the reference region. Regional tau uptake was quantified as volume-weighted average SUVRs across the FreeSurfer regions corresponding to six composite Braak stage regions of interest (ROIs), which reflect the hierarchical spread of tau pathology in Alzheimer’s disease (Gogola et al., 2022): Braak I (entorhinal cortex), Braak II (hippocampus), Braak III (parahippocampal, fusiform, lingual, and amygdala), Braak IV (middle temporal, inferior temporal, temporal pole, cingulate, insula, and thalamus), Braak V (frontal, parietal, precuneus, superior temporal, and lateral occipital regions), and Braak VI (pericalcarine, postcentral, precentral, paracentral, and cuneus regions).

For the primary analyses, participants were classified into two diagnostic groups. Participants with Alzheimer’s disease (AD), early mild cognitive impairment (EMCI), or late mild cognitive impairment (LMCI) were included in the cognitively impaired group (DM), whereas participants classified as cognitively normal (CN) or having significant memory concern (SMC) were included in the cognitively unimpaired group (CU). Hereafter, DM and CU refer to these aggregated analysis groups. To reduce potential confounding between site and diagnostic-group composition, we excluded sites containing participants from only one diagnostic group or fewer than two eligible participants. The resulting analytic sample included 986 participants with complete baseline tau PET measurements. Demographic, genetic, and clinical covariates, however, contained varying degrees of missingness and were addressed using the procedures described in Section 2.2.

### 2.2 Multiple imputation

Missing baseline covariate data were addressed using multiple imputation (MI) under a missing-at-random (MAR) assumption (Rubin, 1976). Imputation was performed using multivariate imputation by chained equations (MICE) (van Buuren & Groothuis-Oudshoorn, 2011) implemented in the mice R package. The imputation model included age, sex, years of education, diagnostic group, race/ethnicity, marital status, and apolipoprotein E (APOE) *ε*4 genotype (0, 1, or 2 alleles). Continuous variables (age, education) were imputed using predictive mean matching (PMM), and categorical variables (diagnostic group, sex, race/ethnicity, marital status, APOE *ε*4) were imputed using logistic or polytomous logistic regression (van Buuren & Groothuis-Oudshoorn, 2011; White et al., 2011). Site was included as a categorical predictor variable in the imputation model to account for potential differences in covariate distributions across sites. Fully observed tau PET outcomes (volume-weighted average SUVRs across Braak regions) were included as auxiliary predictors to improve imputation accuracy. Five imputed datasets (*m* = 5) were generated with 50 iterations. Each completed dataset was subsequently analyzed using the harmonization and statistical procedures described below, with estimates combined across imputations using Rubin’s rules.

### 2.3 Harmonization methods

To reduce residual site-related variability in tau PET SUVR measures, we evaluated four statistical harmonization approaches: two parametric, ComBat with covariate adjustment (ComBat) and ComBat without covariate adjustment (ComBat Null), and two nonparametric, QN and QR. Braak-region SUVRs derived from the ADNI-preprocessed data without additional statistical harmonization (Raw) served as the reference. For all approaches, site was specified as the batch variable. ComBat and ComBat Null were applied jointly across six Braak regions, whereas QN and QR were applied separately to each region. ComBat and QR incorporated age, sex, education, diagnosis, APOE *ε*4, race/ethnicity, and marital status as covariates to preserve biological variation associated with these characteristics, whereas ComBat Null and QN did not incorporate covariate information. Diagnosis was represented using its original five-category classification (AD, EMCI, LMCI, SMC, and CN) during harmonization to preserve biological variability across the spectrum of cognitive impairment.

For the imputed analyses, each harmonization approach was applied separately to each of the five imputed datasets. For methods incorporating covariates (ComBat and QR), the corresponding imputed covariate values from each dataset were used. Downstream statistical analyses were then performed within each harmonized dataset, and estimates were combined across imputations as described in Section 2.4.

#### 2.3.1 ComBat

We denote *Y_ijv_* as the volume-weighted average tau SUVR for subject *i* at site *j* in Braak region *v*. Following Johnson et al. (2007) and its adaptation to neuroimaging data (Fortin et al., 2017, 2018), ComBat models imaging features as:

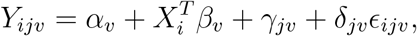

where *α_v_* is the overall mean for region *v*, *X_i_* is the vector of biological covariates with corresponding coefficients *β_v_*, *γ_jv_* is the additive site effect, *δ_jv_* is the multiplicative site effect, and 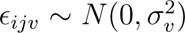. ComBat assumes that, within each site, additive and multiplicative site effects across Braak regions arise from common prior distributions and therefore borrows information across regions to stabilize site-specific parameter estimates, particularly when site sample sizes are small (Beer et al., 2020; Fortin et al., 2018; Johnson et al., 2007). The resulting empirical Bayes estimates 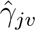 and 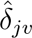 are used to compute the harmonized values as:

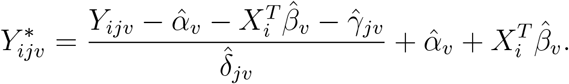

#### 2.3.2 Quantile normalization (QN)

QN harmonizes each Braak region by mapping site-specific empirical distributions to a common reference distribution (Bolstad et al., 2003; Zhang et al., 2026). For each Braak region *v*, let 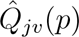 denote the empirical quantile function for site *j* at quantile level *p* ∈ [0, 1]. The common reference distribution is defined by averaging the site-specific quantiles,

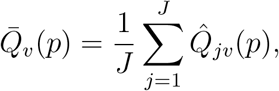

where *J* is the total number of sites. When site sample sizes differ, site-specific quantiles are interpolated onto a common quantile grid before averaging. Each observed SUVR value is then mapped to the corresponding quantile of the reference distribution.

This procedure aligns the empirical distributions across sites for each Braak region, reducing systematic differences in location, scale, and shape. QN was applied separately to each of the six Braak composite regions using the batchtma R package (Stopsack et al., 2021).

#### 2.3.3 Quantile regression (QR)

QR harmonization assumes that, after accounting for covariates, sites with similar participant characteristics would share the same quantiles in the absence of site effects (Bann et al., 2020). Building on QR (Cade & Noon, 2003; R. Koenker & Hallock, 2001), this approach models the 25th and 75th percentiles separately, allowing covariate effects to differ across quantiles and site-related differences in location and interquartile range to be addressed.

For each Braak region *v*, we fitted two quantile regression models at *τ* = 0.25 and *τ* = 0.75 respectively, using a modified Barrodale–Roberts simplex algorithm (R. W. Koenker & D’Orey, 1987). The model for the *τ* -th conditional quantile is given by:

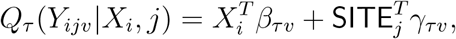

where *X_i_* is the vector of covariates described above; *β_τv_* captures covariate effects at quantile *τ* ; and *γ_τv_* represents site-specific deviations. From these models, we extracted three quantities per site and region: 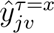, the empirical *x*-th quantile for site *j*; 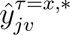, the covariate-adjusted *x*-th quantile for site *j*; 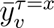, the overall *x*-th quantile pooled across all sites.

We implemented QR using the batchtma R package (Stopsack et al., 2021), with the same covariates used in ComBat. The harmonized SUVR for each subject is computed as:

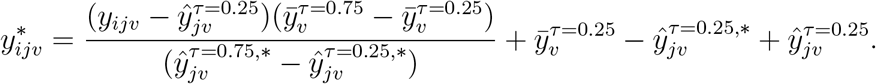

This location-scale transformation uses the site-specific 25th percentile as the location offset and the interquartile range (IQR; 75th–25th percentile) as the scale factor, removing both additive and multiplicative site effects while preserving covariate associations.

### 2.4 Statistical analysis

To evaluate the combined impact of MI and harmonization, we compared CC and imputed analyses along two complementary dimensions: 1) harmonization effectiveness, characterized by residual site-related variability after statistical harmonization using three metrics; and 2) preservation of biological effects, characterized by differences in tau SUVR between DM and CU participants using effect sizes and regression coefficients.

To estimate disease-group differences while accounting for demographic, genetic, and clinical characteristics, we fitted a separate linear regression model for each Braak region *v*:

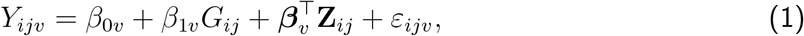

where *i*, *j*, and *v* index subjects, sites, and Braak regions, respectively, as defined in Section 2.3. *G_ij_* ∈ {0, 1} is a binary indicator of disease group (1 = DM; 0 = CU), and the vector **Z***_ij_* represents covariates (age, sex, education, race/ethnicity, marital status, and APOE *ε*4), with corresponding region-specific coefficient vector ***β****_v_* . The error terms were assumed to be independent and normally distributed, 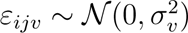.

For the imputed analyses, the model was fitted separately within each imputed dataset, and the resulting estimates and their variances were combined using Rubin’s rules (Rubin, 1987). Confidence intervals (CIs) and *p*-values were calculated based on the combined estimates and corresponding standard errors (SEs), using a *t*-distribution with Barnard–Rubin-adjusted degrees of freedom (Barnard & Rubin, 1999). The precision of imputed results relative to the CC analysis was assessed using the ratio SE_MI_*/*SE_CC_, with values below 1 indicating smaller SEs in the imputed analyses. The fraction of missing information (FMI) was used to quantify uncertainty attributable to missing data. Lower FMI values indicate a smaller contribution of missingness to the total variance (White et al., 2011).

#### 2.4.1 Assessment of harmonization effectiveness

Harmonization effectiveness was assessed using three metrics, computed separately for each Braak region and harmonization method for the CC and imputed analyses:

(1). **Pairwise site mean and SD differences.** Absolute differences in site-level mean SUVRs and SDs were calculated between all pairs of sites, with lower values indicating greater homogeneity across sites. For the MI analysis, pairwise differences were calculated separately within each imputed dataset and then averaged across the five imputations.
(2). **Covariate-adjusted intraclass correlation coefficients.** ICCs quantify the proportion of residual variance attributable to site effects after adjusting for demographic and biological covariates (Shrout & Fleiss, 1979). Using residuals from Eq. (1), the ICC was calculated for each Braak region as:

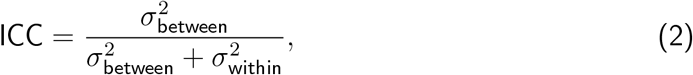

where 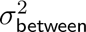 is the variance of site-specific mean residuals and 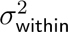 is the pooled within-site variance (weighted by site sample sizes). For MI analyses, estimates were calculated separately within each imputed dataset and averaged across the five imputations. For CC analyses, a single estimate was calculated directly.

#### 2.4.2 Assessment of biological effects

To assess the preservation of biological effects after statistical harmonization, we used three complementary metrics to evaluate the stability of statistical inference across harmonization methods, and compared the results between CC and imputed datasets.

(1). **Standardized effect sizes.** For each harmonization method and Braak region, we calculated Hedges’ *g* (Cohen, 1988), a bias-corrected standardized mean difference between DM and CU groups. For imputed analyses, Hedges’ *g* and its sampling variance, estimated using a large-sample approximation, were calculated separately within each imputed dataset and combined according to Rubin’s rules (Rubin, 1987). Pooled 95% CIs were calculated using a *t*-distribution with Barnard-Rubin-adjusted degrees of freedom (Barnard & Rubin, 1999).
(2). **Regression coefficients and statistical significance.** From Eq. (1), *β*_1_ represents the covariate-adjusted mean SUVR difference between the DM and CU groups. For the CC analysis, 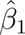, its 95% CI, and the corresponding *p*-value were obtained from ordinary least squares regression. For MI analyses, coefficient estimates and their estimated variances were obtained within each imputed dataset and combined using Rubin’s rules (Rubin, 1987), accounting for within- and between-imputation variability. Pooled 95% CIs and two-sided *p*-values were calculated using a *t*-distribution with Barnard–Rubin-adjusted degrees of freedom (Barnard & Rubin, 1999). Statistical significance was assessed for *H*_0_ : *β*_1_ = 0 at *α* = 0.05. We reported − log_10_(*p*) values to compare statistical evidence across methods, with higher values indicating stronger evidence against the null.

As a sensitivity analysis, we additionally evaluated meta-temporal SUVR, an established composite measure used in tau PET studies (Gogola et al., 2023; Jack et al., 2017), using the same imputation, harmonization, and evaluation procedures. For ComBat and ComBat Null, meta-temporal SUVR was included alongside six Braak composites as a seventh imaging feature, and only the meta-temporal results from this analysis were reported. Details are provided in the Supplementary Materials.

## 3 Results

### 3.1 Study population and missingness

Baseline demographic and clinical characteristics for the full analytic sample before imputation (*n* = 986) are presented in Table 1. The sample had a mean age of 71.0 years (*SD* = 7.06), a mean of 16.4 years of education (*SD* = 2.45), and was predominantly white (72.1%). Among participants with observed baseline diagnosis, 69 had AD, 170 had EMCI, 161 had LMCI, 265 had SMC, and 236 were CN.

**Table 1:**
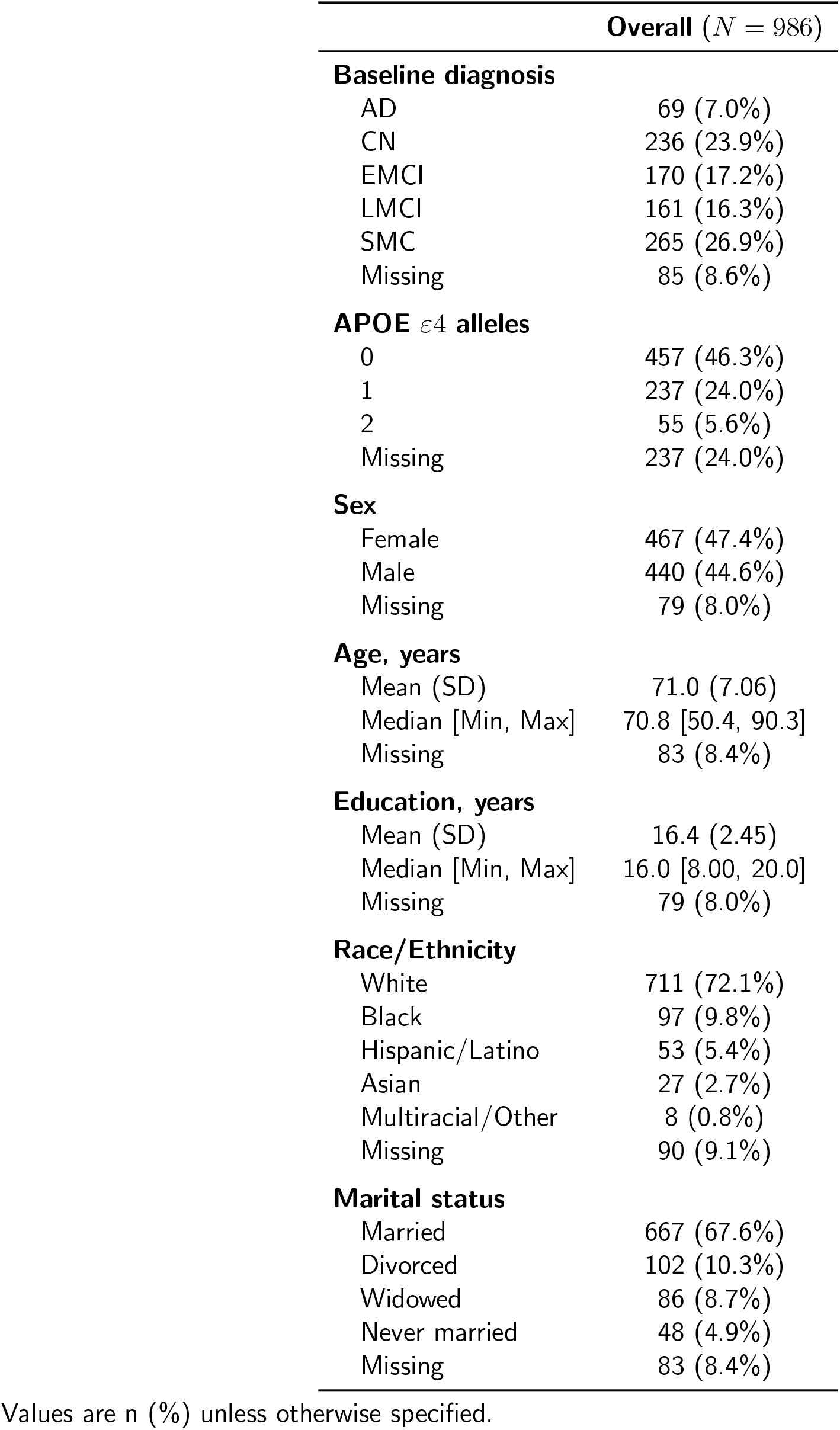
Overall sample characteristics and data completeness.

|  | **Overall** (*N* = 986) |
|---|---|
| **Baseline diagnosis** |  |
| AD | 69 (7.0%) |
| CN | 236 (23.9%) |
| EMCI | 170 (17.2%) |
| LMCI | 161 (16.3%) |
| SMC | 265 (26.9%) |
| Missing | 85 (8.6%) |
| **APOE** *ε*4 **alleles** |  |
| 0 | 457 (46.3%) |
| 1 | 237 (24.0%) |
| 2 | 55 (5.6%) |
| Missing | 237 (24.0%) |
| **Sex** |  |
| Female | 467 (47.4%) |
| Male | 440 (44.6%) |
| Missing | 79 (8.0%) |
| **Age, years** |  |
| Mean (SD) | 71.0 (7.06) |
| Median [Min, Max] | 70.8 [50.4, 90.3] |
| Missing | 83 (8.4%) |
| **Education, years** |  |
| Mean (SD) | 16.4 (2.45) |
| Median [Min, Max] | 16.0 [8.00, 20.0] |
| Missing | 79 (8.0%) |
| **Race/Ethnicity** |  |
| White | 711 (72.1%) |
| Black | 97 (9.8%) |
| Hispanic/Latino | 53 (5.4%) |
| Asian | 27 (2.7%) |
| Multiracial/Other | 8 (0.8%) |
| Missing | 90 (9.1%) |
| **Marital status** |  |
| Married | 667 (67.6%) |
| Divorced | 102 (10.3%) |
| Widowed | 86 (8.7%) |
| Never married | 48 (4.9%) |
| Missing | 83 (8.4%) |

Values are n (%) unless otherwise specified.

Missingness was highest for the APOE *ε*4 genotype (24.0%); missingness in the remaining covariates ranged from 8.0% to 9.1%, including age (8.4%), sex (8.0%), education (8.0%), baseline diagnosis (8.6%), race/ethnicity (9.1%), and marital status (8.4%). Baseline tau PET SUVR outcomes were fully observed in the analytic sample. Supplementary Table S1 provides detailed characteristics for the CC sample (*n* = 710).

### 3.2 Harmonization effectiveness

#### 3.2.1 Site-level mean and variance homogeneity

Pairwise site differences in means and SDs are shown in Figure 1. Full boxplots for all imputed datasets across harmonization methods and Braak regions are provided in Supplementary Figure S2.

**Figure 1:**
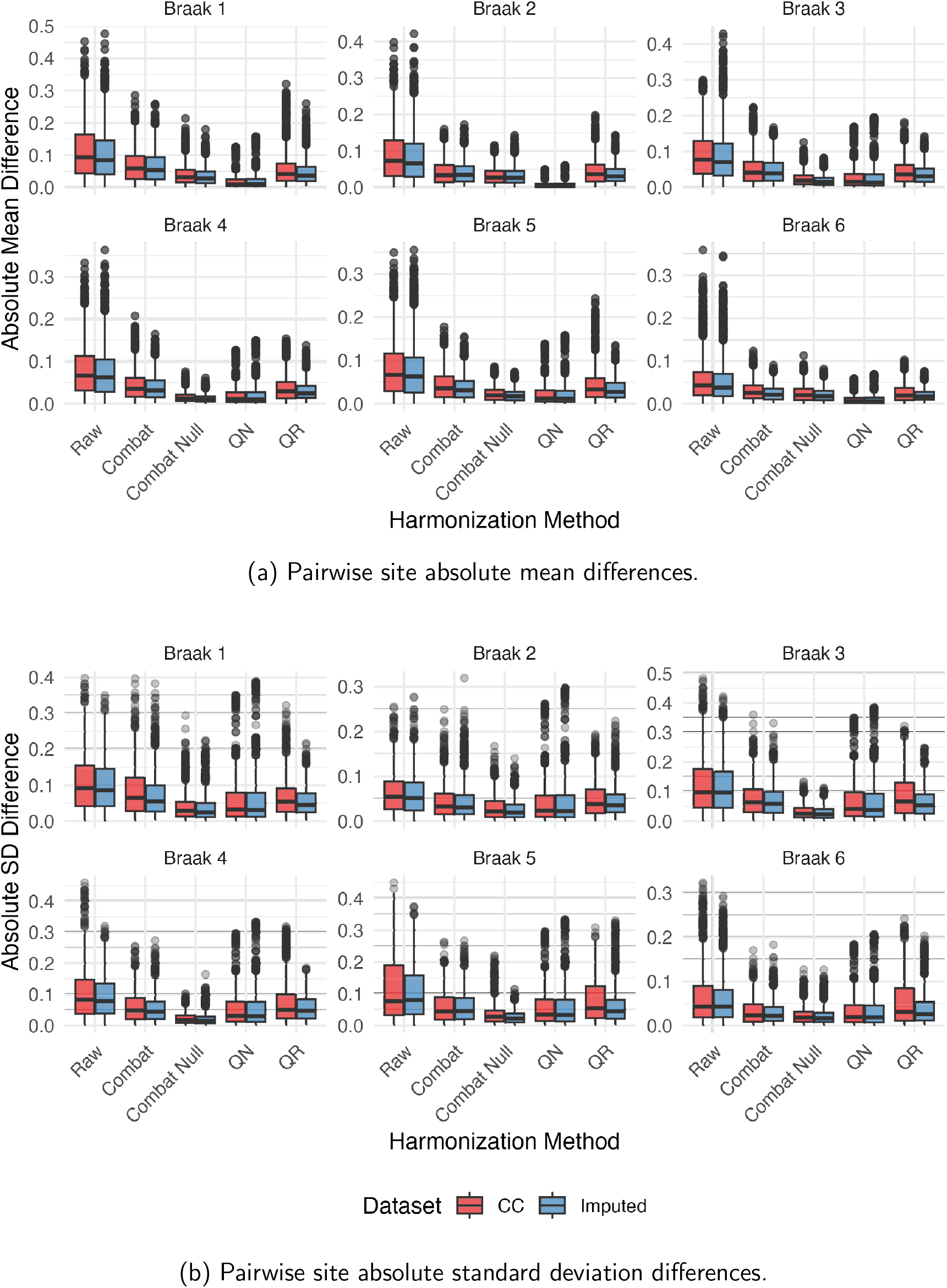
Site-level consistency metrics comparing complete case (red) and imputed (blue) analyses across harmonization methods and Braak regions. (a) Distribution of absolute differences in mean tau SUVR between all site pairs. (b) Distribution of absolute differences in within-site standard deviations between all site pairs. Lower values indicate better harmonization consistency.

Raw data exhibited substantial between-site heterogeneity in both mean and variance across all Braak regions, with pairwise mean differences showing outliers extending to 0.3 − 0.5 SUVR units. All harmonization methods reduced these differences. For mean differences, ComBat Null and QN produced the most compact distributions, while QR showed slightly greater residual site-level variability in several regions. A similar pattern was observed for within-site SD differences: ComBat Null consistently achieved the greatest reduction across Braak regions, followed by QN and ComBat, while QR showed higher dispersion. Overall patterns were consistent between CC and imputed analyses, with similar method rankings and comparable distributions observed across all Braak regions.

#### 3.2.2 Site variance reduction using covariate-adjusted ICC

Covariate-adjusted ICCs are displayed in Figure 2. Raw data exhibited the highest values across all Braak regions, indicating substantial residual between-site variability after adjustment for demographic and clinical covariates. All harmonization methods markedly reduced ICCs, indicating substantial reductions in residual site-related variability. ComBat achieved the lowest values in most Braak regions, whereas QN yielded the lowest values in Braak regions II and VI. ComBat Null and QR generally showed slightly higher ICCs. The overall pattern across methods and Braak regions was similar between CC and imputed analyses. Detailed estimates are provided in the Supplementary Table S2.

**Figure 2:**
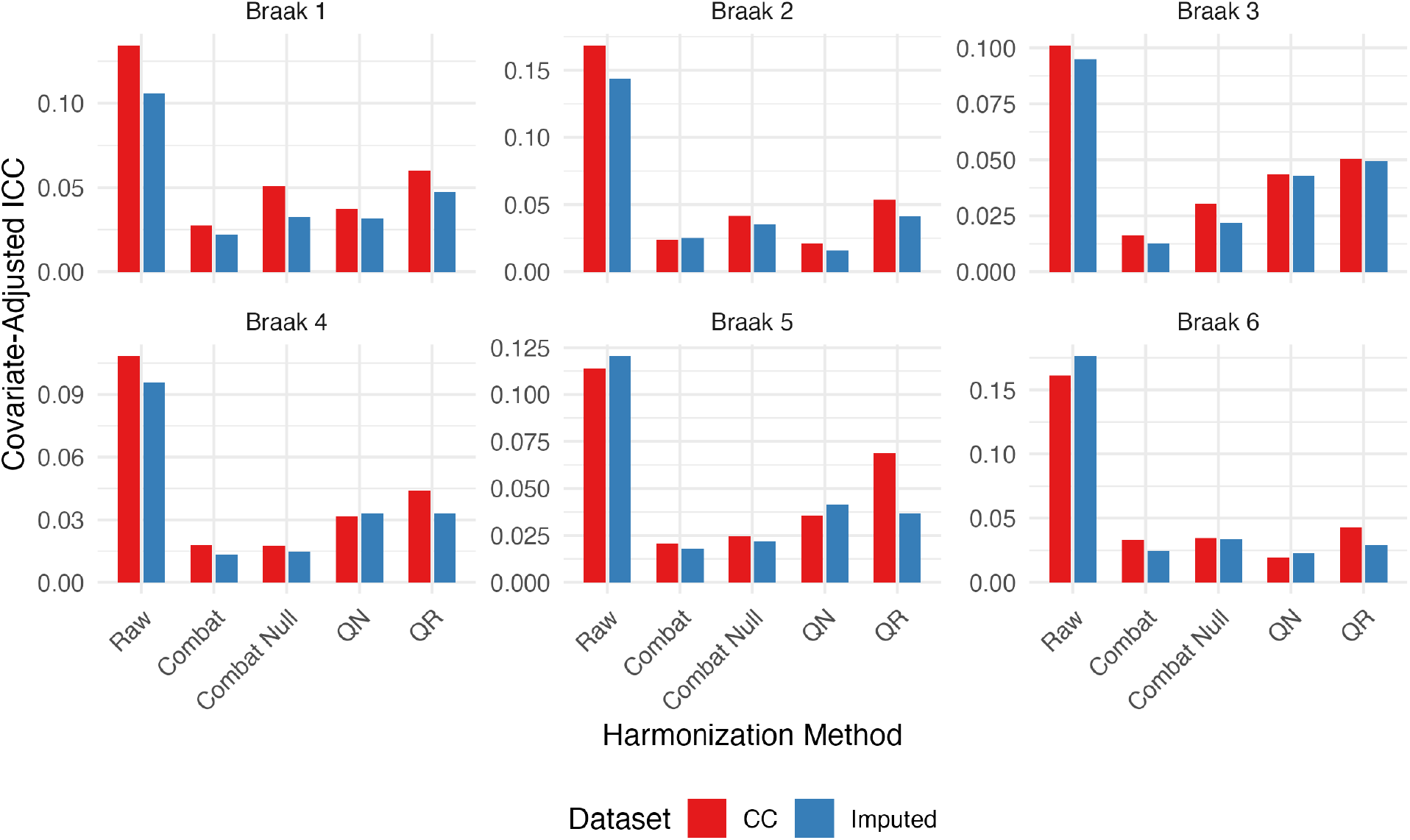
Intraclass correlation coefficients comparing complete case (red) and imputed (blue) analyses across harmonization methods and Braak regions. Approximate ICCs of site calculated from residualized SUVR after controlling for disease group (DM versus CU), age, sex, education, APOE *ε*4, race/ethnicity, and marital status. Lower values indicate less dispersion of site mean residuals relative to within-site residual variability.

### 3.3 Preservation of biological effects

#### 3.3.1 Standardized effect sizes

Table 2 presents Hedges’ *g*, a standardized measure of tau SUVR differences between DM and CU groups across harmonization methods and Braak regions.

**Table 2:**
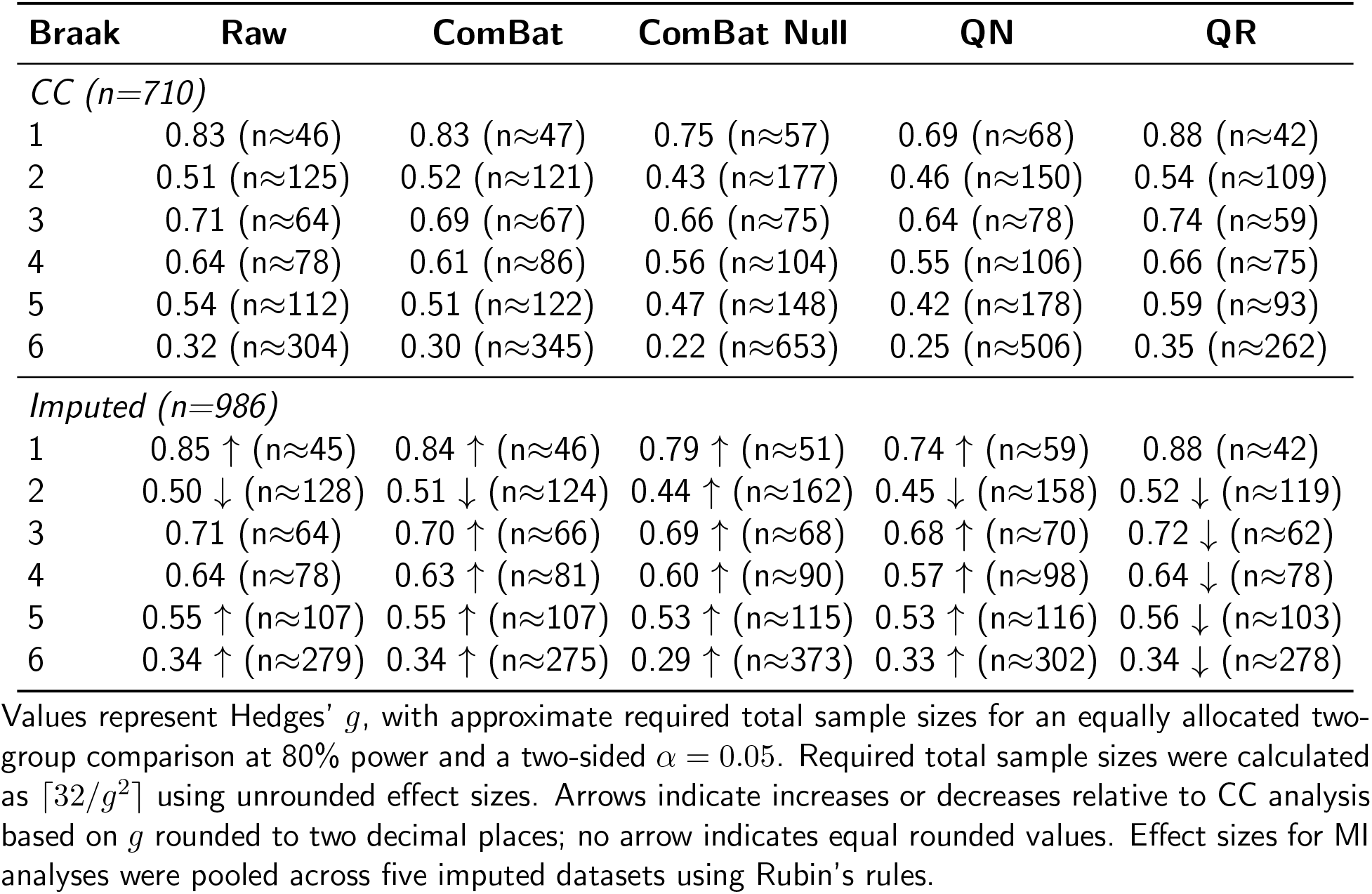
Hedges’ *g* (required sample sizes) for DM vs CU group discrimination comparing CC vs imputed analyses.

| **Braak** | **Raw** | **ComBat** | **ComBat Null** | **QN** | **QR** |
|---|---|---|---|---|---|
| *CC (n=710)* |  |  |  |  |  |
| 1 | 0.83 (n≈46) | 0.83 (n≈47) | 0.75 (n≈57) | 0.69 (n≈68) | 0.88 (n≈42) |
| 2 | 0.51 (n≈125) | 0.52 (n≈121) | 0.43 (n≈177) | 0.46 (n≈150) | 0.54 (n≈109) |
| 3 | 0.71 (n≈64) | 0.69 (n≈67) | 0.66 (n≈75) | 0.64 (n≈78) | 0.74 (n≈59) |
| 4 | 0.64 (n≈78) | 0.61 (n≈86) | 0.56 (n≈104) | 0.55 (n≈106) | 0.66 (n≈75) |
| 5 | 0.54 (n≈112) | 0.51 (n≈122) | 0.47 (n≈148) | 0.42 (n≈178) | 0.59 (n≈93) |
| 6 | 0.32 (n≈304) | 0.30 (n≈345) | 0.22 (n≈653) | 0.25 (n≈506) | 0.35 (n≈262) |
| *Imputed (n=986)* |  |  |  |  |  |
| 1 | 0.85 ↑ (n≈45) | 0.84 ↑ (n≈46) | 0.79 ↑ (n≈51) | 0.74 ↑ (n≈59) | 0.88 (n≈42) |
| 2 | 0.50 ↓ (n≈128) | 0.51 ↓ (n≈124) | 0.44 ↑ (n≈162) | 0.45 ↓ (n≈158) | 0.52 ↓ (n≈119) |
| 3 | 0.71 (n≈64) | 0.70 ↑ (n≈66) | 0.69 ↑ (n≈68) | 0.68 ↑ (n≈70) | 0.72 ↓ (n≈62) |
| 4 | 0.64 (n≈78) | 0.63 ↑ (n≈81) | 0.60 ↑ (n≈90) | 0.57 ↑ (n≈98) | 0.64 ↓ (n≈78) |
| 5 | 0.55 ↑ (n≈107) | 0.55 ↑ (n≈107) | 0.53 ↑ (n≈115) | 0.53 ↑ (n≈116) | 0.56 ↓ (n≈103) |
| 6 | 0.34 ↑ (n≈279) | 0.34 ↑ (n≈275) | 0.29 ↑ (n≈373) | 0.33 ↑ (n≈302) | 0.34 ↓ (n≈278) |

Values represent Hedges’ *g*, with approximate required total sample sizes for an equally allocated two-group comparison at 80% power and a two-sided $\alpha = 0.05$. Required total sample sizes were calculated as $\lceil 32/g^2 \rceil$ using unrounded effect sizes. Arrows indicate increases or decreases relative to CC analysis based on *g* rounded to two decimal places; no arrow indicates equal rounded values. Effect sizes for MI analyses were pooled across five imputed datasets using Rubin’s rules.

In the CC analysis, effect sizes ranged from 0.22 to 0.88 across methods and Braak regions, spanning small to large effects by conventional thresholds (Cohen, 1988). Raw data showed effect sizes ranging from 0.32 (Braak VI) to 0.83 (Braak I). Across harmonization methods, ComBat produced effect size estimates nearly identical to those from raw data (0.30–0.83). ComBat Null and QN showed modest reductions in later Braak regions, particularly in Braak VI (0.22 and 0.25, respectively). In contrast, QR consistently yielded the largest effect sizes across all Braak regions in the CC analysis (0.35–0.88), corresponding to the smallest approximate required total sample sizes (42–262 participants). This suggests slightly improved group discrimination performance under QR.

In imputed analyses, effect sizes ranged from 0.29 to 0.88 and followed the same spatial pattern observed in the CC analysis, with the largest effects in Braak I and the smallest in Braak VI. All harmonization methods preserved substantial effect sizes relative to raw data (0.34–0.85) and demonstrated broadly similar relative performance to that in the CC analysis. ComBat maintained effect sizes closest to raw values (0.34–0.84), followed by ComBat Null (0.29–0.79), while QN showed slightly lower effect sizes (0.33–0.74). QR yielded effect sizes comparable to or slightly above raw values (0.34–0.88). Pooled SEs ranged from 0.067 to 0.071, and SE ratios were consistently below 1 (median: 0.89), suggesting improved precision with imputation. FMI values were generally low to moderate (0.04–0.14), indicating modest uncertainty attributable to missing data. All group differences remained highly significant (*p <* 0.001) across methods and Braak regions. Detailed estimates across methods, Braak regions, and datasets are reported in Supplementary Table S3.

Overall, effect size estimates were highly consistent between CC and imputed analyses. The absolute differences in Hedges’ *g* were small (mean |Δ*g*| = 0.027, *SD* = 0.025), with 25 out of 30 comparisons showing changes smaller than 0.05. Percentage changes ranged from –4.96% to 32.40%, with 25 out of 30 comparisons within ±10%. In terms of direction, effect sizes increased after imputation in 21 of 30 comparisons. Increases were observed for nearly all regions under Raw, ComBat, ComBat Null, and QN, with the largest increases for ComBat Null and QN in Braak V and VI, whereas QR showed small decreases in five of six regions. Method-specific mean differences were small: Raw (0.006), ComBat (0.016), ComBat Null (0.044), QN (0.046), and QR (-0.015).

#### 3.3.2 Regression coefficients and statistical significance

Figure 3 shows the regression-based estimates of disease group differences (*β*_1_*_v_* from Eq. 1) across harmonization methods and Braak regions, while Figure 4 shows the corresponding − log_10_(*p*) values, where higher values indicate stronger evidence against the null hypothesis of no group difference.

**Figure 3:**
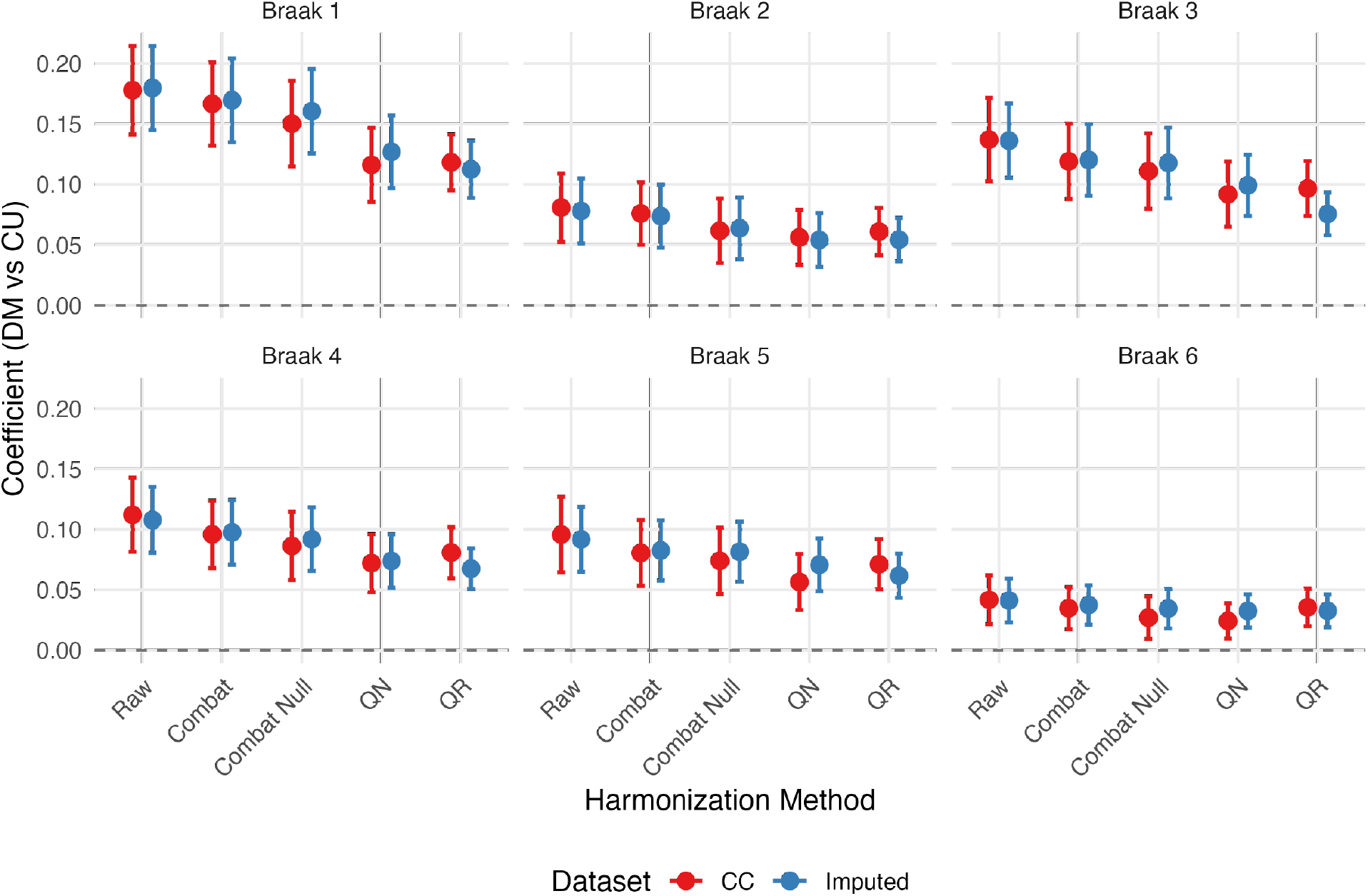
Group effect estimates comparing CC (red) and imputed (blue) analyses. Points show regression coefficients (DM vs CU) with 95% CIs. Estimates from imputed analyses were pooled using Rubin’s rules.

**Figure 4:**
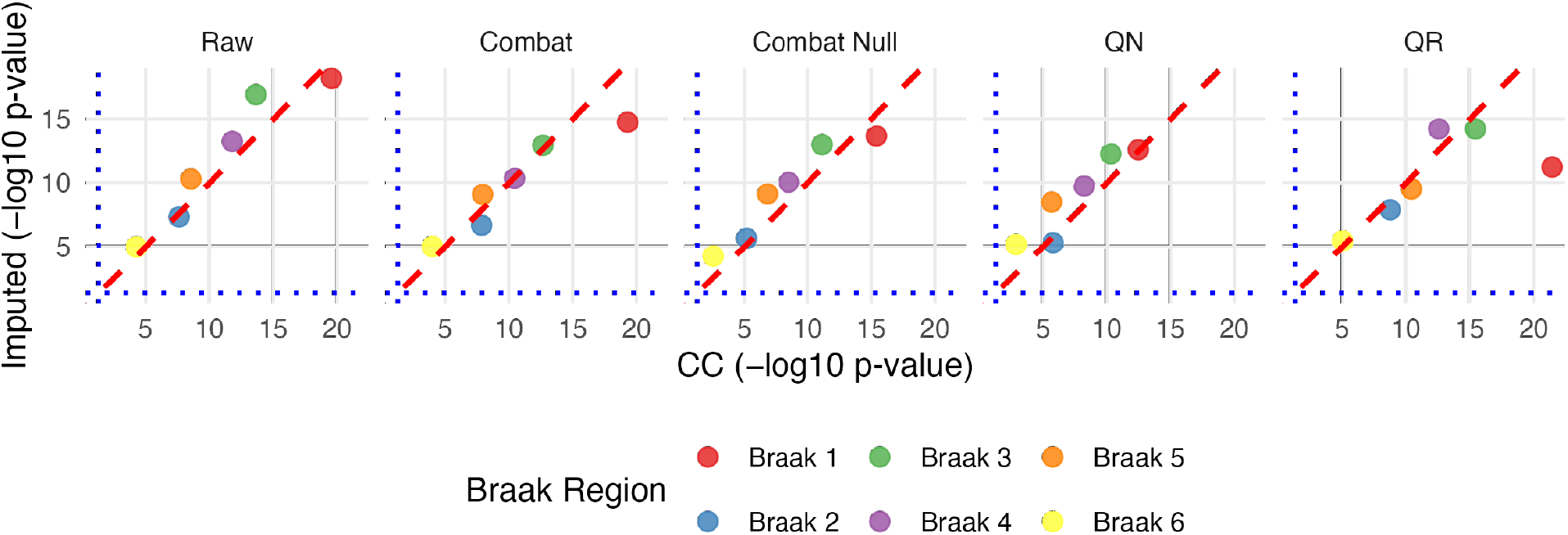
Statistical significance between CC and imputed analyses. Scatter plots show − log_10_(*p*) values for group effect comparisons. The red dashed diagonal indicates perfect agreement; blue dotted lines show the *p* = 0.05 significance threshold. For imputed analyses, estimates and corresponding inference were pooled across the five imputations using Rubin’s rules.

For the CC analysis, raw data consistently produced the largest regression coefficients across all Braak regions, reflecting the largest covariate-adjusted group separation. All harmonization methods yielded slightly reduced coefficients relative to raw data. Among the harmonized datasets, ComBat and ComBat Null generally maintained relatively higher coefficients, whereas QN and QR showed more noticeable reductions across regions. All disease group comparisons were statistically significant (*p <* 0.05) across harmonization methods and Braak regions. QR exhibited the strongest statistical evidence, followed by raw data and ComBat, whereas ComBat Null showed slightly reduced statistical strength and QN showed a more pronounced reduction. Across all methods, statistical evidence was strongest in Braak I and weakest in Braak VI, with the overall ordering approximately Braak I *>* III *>* IV *>* V *>* II *>* VI, consistent with the regional pattern of effect sizes in Section 3.3.1. CIs were narrower in later Braak regions, particularly in Braak VI.

For imputed analyses, regression coefficients followed similar patterns. Harmonization resulted in modest reductions in regression coefficients compared with raw data. Among the harmonized datasets, ComBat generally yielded the largest coefficients, followed by ComBat Null, while QN and QR showed modest reductions. Statistical significance patterns were also broadly similar to those in the CC analysis, and all disease group comparisons remained statistically significant across methods and Braak regions. Raw data showed the strongest statistical evidence, with QR performing comparably in most regions except for a notable reduction in Braak I. Other harmonization methods, particularly QN, showed modest reductions in statistical strength. CIs were generally narrower in the imputed analyses. SE ratios ranged from 0.77 to 0.99 (median: 0.93), corresponding to 1–23% reductions in SEs. FMI values ranged from 0.06 to 0.31 (median: 0.16), indicating modest imputation uncertainty appropriately incorporated through Rubin’s rules.

Comparing CC and imputed analyses, regression coefficients were highly concordant across harmonization methods and regions. The direction of changes between CC and imputed estimates varied by method and Braak region. Notably, QR produced lower coefficient estimates in the imputed analyses compared with CC across all regions. Differences in coefficient estimates were generally small, and CIs were generally narrower in the imputed analyses, consistent with the observed gains in precision. Similarly, most points in Figure 4 remained on or slightly above the diagonal reference line, indicating similar or modestly stronger statistical evidence after imputation, particularly in later Braak regions. Some points fell below the diagonal for specific method–region combinations, particularly QR in Braak I, consistent with the decreases in regression coefficient estimates under imputation. Overall, the method-specific and regional patterns were highly consistent between the CC and imputed analyses. Detailed regression coefficients, CIs, SE ratios, and FMI values are provided in Supplementary Table S4, and detailed *p*-values are provided in Supplementary Table S5.

In the meta-temporal SUVR sensitivity analysis, harmonization reduced site variability while retaining DM–CU differences, and imputed analyses yielded smaller standard errors than complete-case analyses. Detailed results are provided in Supplementary Table S6.

## 4 Discussion

This study evaluated the joint impact of MI and several harmonization approaches, including ComBat with and without covariates, QN, and QR, on multisite [^18^F]flortaucipir tau PET data from ADNI. Statistical harmonization was applied to ADNI-preprocessed baseline SUVRs across six Braak composite regions, and MI was performed using MICE with *m* = 5 imputations. The results demonstrated that all harmonization methods substantially reduced residual site-related variability compared with raw data, although their relative performance differed by evaluation metric. Importantly, MI did not compromise harmonization effectiveness: harmonization methods showed broadly similar patterns of reduction in site heterogeneity in the CC and imputed analyses, with heterogeneity metrics generally comparable or lower after MI.

Furthermore, disease group differences were preserved after statistical harmonization and imputation. In regression models adjusted for age, sex, education, race/ethnicity, APOE *ε*4, and marital status, disease group effects (DM vs CU) followed a consistent regional pattern, with the strongest group differences in Braak I and the weakest in Braak VI. Compared with the CC analysis, the imputed analyses generally yielded slightly larger effect sizes and regression coefficients, accompanied by narrower CIs and smaller SEs. These findings suggested that imputation preserved disease-related effects with slightly improved statistical efficiency by retaining more participants. The sensitivity analysis of meta-temporal SUVR, a composite widely used in clinical tau PET studies (Gogola et al., 2023; Jack et al., 2017), yielded consistent results, indicating that the observed harmonization and imputation patterns were not specific to the Braak regions. Taken together, the results supported the combined use of MI and harmonization in multisite tau PET studies with incomplete covariate data. Beyond the performance of individual harmonization methods, this study extended existing harmonization research by evaluating multiple imputation and harmonization within the same multisite tau PET framework.

Our findings aligned with previous neuroimaging harmonization studies demonstrating that ComBat markedly reduced batch effects in diffusion and structural MRI (Fortin et al., 2017; Pomponio et al., 2020). In our analysis, covariate-adjusted ComBat yielded the lowest observed approximate ICC values in most Braak regions, whereas ComBat Null produced smaller pairwise SD differences overall. This dissociation paralleled the findings of (Yang et al., 2024), who observed that ComBat Null was the most effective at improving inter-tracer agreement in amyloid PET, while covariate-adjusted variants better preserved biological covariates.

Quantile-based methods also reduced site-level variability, extending approaches used in tissue microarray biomarker studies (Stopsack et al., 2021) to PET imaging contexts. However, the stronger distributional alignment achieved by QN was accompanied by greater attenuation of effect sizes, suggesting potential over-harmonization. This aggressive distributional alignment might have removed some biologically relevant between-site heterogeneity (Chen et al., 2022). In contrast, QR adjusted selected conditional quantiles while accounting for covariates, yielding the largest standardized effect sizes in the CC analysis but reduced regression coefficients relative to raw data. QR was also the only method whose estimates consistently decreased after imputation. These findings suggested that distribution-based harmonization approaches might differ in their ability to balance site-effect reduction and preservation of biological variation.

Missing data remain a persistent challenge in multisite neuroimaging studies, particularly in large observational cohorts such as ADNI. In our study, more than one quarter of participants were excluded from the CC analysis, primarily due to missing APOE *ε*4 genotype and demographic covariates. This pattern was consistent with the broader ADNI literature, in which missingness varied between biomarkers and clinical groups (Lo et al., 2012). Such exclusions could reduce statistical efficiency and introduce bias when missingness was associated with disease severity or other covariates. MI retained these participants and led to modestly increased disease group effect estimates and reduced SEs across most harmonization settings. The observed precision gains were broadly consistent with expectations based on the increase in sample size 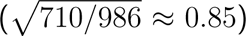.

FMI values were generally low to moderate, indicating that missingness accounted for a limited to moderate proportion of total variance (White et al., 2011). Importantly, site heterogeneity metrics showed consistent patterns between the CC and imputed analyses, suggesting that MI did not destabilize harmonization performance.

Recent methodological studies in ADNI showed that MICE performed favorably compared with alternative imputation approaches in predictive modeling and longitudinal disease progression analyses (Aracri et al., 2025; Chandrasekaran et al., 2024). Similarly, machine-learning-based imputation methods such as missForest were evaluated for handling incomplete clinical and neuroimaging data (Aracri et al., 2023). In the context of harmonization, MICE has also been used as a preprocessing step before ComBat harmonization, for example, to impute missing regional diffusion MRI measures (Alasmar et al., 2026). However, evidence on how MI interacted with multisite harmonization procedures and affected downstream statistical inference remained limited. Our findings addressed this gap by demonstrating that MICE could be effectively combined with harmonization approaches in cross-sectional tau PET analyses without compromising harmonization effectiveness or the preservation of biological group differences.

Several limitations should be acknowledged. First, the ADNI cohort was predominantly white and highly educated, which may limit generalizability to more diverse populations. Second, only a subset of harmonization strategies was examined; newer extensions such as ComBat-GAM (Pomponio et al., 2020), CovBat (Chen et al., 2022), ComBatLS (Gardner et al., 2025), and deep learning methods (An et al., 2025; Dewey et al., 2019; Hu et al., 2024) may provide additional flexibility. Third, our analysis was limited to [^18^F]flortaucipir, and the findings may not generalize to other tau PET tracers with different binding and off-target characteristics. Future studies should evaluate whether similar harmonization and imputation patterns are observed with newer-generation tau tracers. Fourth, our analysis focused only on cross-sectional baseline data. Future work should extend this framework to longitudinal tau PET trajectories using approaches such as longitudinal ComBat (Beer et al., 2020) and joint longitudinal imputation models that account for the nested structure of repeated measures within subjects within sites. Developing unified frameworks that simultaneously handle site effects and missing data would streamline analysis pipelines.

In summary, our findings support the integration of multiple imputation with harmonization as a practical approach for analyzing multisite neuroimaging data with incomplete covariate information. This approach enables the inclusion of additional participants and improves the precision of downstream estimates while maintaining harmonization effectiveness and preserving biological associations.

## Declaration of the use of AI

AI tools were not used to design or conduct the research, perform statistical analyses, or generate the study results reported in this manuscript. During manuscript preparation, the authors used ChatGPT and Claude to assist with rewording draft text, grammar, and improvements to clarity and readability. All AI-assisted text was reviewed, verified, and edited by the authors, who take full responsibility for the content of the manuscript.

## Data and Code Availability

Data used in this study were obtained from the ADNI and are available to qualified investigators through the Image and Data Archive (IDA) at the Laboratory of Neuro Imaging (LOIN), subject to approval and compliance with the ADNI Data Use Agreement. Participant-level ADNI data and derived participant-level datasets cannot be redistributed by the authors. Analysis code used for multiple imputation, harmonization, statistical analyses, and figure generation is available upon request.

## Author Contributions

Y.R.: Conceptualization, methodology, software, data curation, formal analysis, visualization, writing–original draft, and writing–review & editing. Y.D.: Methodology and writing–review & editing. D.M.: Data curation, methodology, and writing–review & editing. A.D.C.: Writing– review & editing. Q.W.: Conceptualization, methodology, supervision, and writing–review & editing. D.L.T.: Conceptualization, methodology, supervision, project administration, funding acquisition, and writing–review & editing.

## Acknowledgements

Data collection and sharing for the Alzheimer’s Disease Neuroimaging Initiative (ADNI) is funded by the National Institute on Aging (National Institutes of Health Grant U19 AG024904). The grantee organization is the Northern California Institute for Research and Education. In the past, ADNI has also received funding from the National Institute of Biomedical Imaging and Bioengineering, the Canadian Institutes of Health Research, and private sector contributions through the Foundation for the National Institutes of Health (FNIH) including generous contributions from the following: AbbVie, Alzheimer’s Association; Alzheimer’s Drug Discovery Foundation; Araclon Biotech; BioClinica, Inc.; Biogen; Bristol-Myers Squibb Company; CereSpir, Inc.; Cogstate; Eisai Inc.; Elan Pharmaceuticals, Inc.; Eli Lilly and Company; EuroImmun; F. Hoffmann-La Roche Ltd and its affiliated company Genentech, Inc.; Fujirebio; GE Healthcare; IXICO Ltd.; Janssen Alzheimer Immunotherapy Research & Development, LLC.; Johnson & Johnson Pharmaceutical Research &D evelopment LLC.; Lumosity; Lundbeck; Merck & Co., Inc.; Meso Scale Diagnostics, LLC.; NeuroRx Research; Neurotrack Technologies; Novartis Pharmaceuticals Corporation; Pfizer Inc.; Piramal Imaging; Servier; Takeda Pharmaceutical Company; and Transition Therapeutics.

## Supplementary Materials

Supplementary material for this article is available.

