## Supplementary Materials for "Integrating multiple imputation with harmonization in multisite neuroimaging data: implications for site effects and biological inference"

### S1 Supplementary Tables

**Table S1:** Baseline characteristics by diagnostic group (CC analysis, n=710)

| | AD (N=55) | CN (N=202) | EMCI (N=138) | LMCI (N=117) | SMC (N=198) | Overall (N=710) |
|---|---|---|---|---|---|---|
| **APOE $\varepsilon 4$ alleles** | | | | | | |
| 0 | 16 (29.1%) | 146 (72.3%) | 79 (57.2%) | 64 (54.7%) | 127 (64.1%) | 432 (60.8%) |
| 1 | 25 (45.5%) | 52 (25.7%) | 45 (32.6%) | 39 (33.3%) | 65 (32.8%) | 226 (31.8%) |
| 2 | 14 (25.5%) | 4 (2.0%) | 14 (10.1%) | 14 (12.0%) | 6 (3.0%) | 52 (7.3%) |
| **Sex** | | | | | | |
| Female | 20 (36.4%) | 114 (56.4%) | 51 (37.0%) | 48 (41.0%) | 125 (63.1%) | 358 (50.4%) |
| Male | 35 (63.6%) | 88 (43.6%) | 87 (63.0%) | 69 (59.0%) | 73 (36.9%) | 352 (49.6%) |
| **Age, years** | | | | | | |
| Mean (SD) | 74.0 (8.39) | 71.2 (6.01) | 71.0 (7.42) | 71.6 (7.79) | 71.4 (6.20) | 71.5 (6.88) |
| Median [Min, Max] | 74.8 [55.3, 89.2] | 70.8 [55.1, 86.3] | 72.5 [55.0, 86.4] | 71.9 [55.0, 90.0] | 70.1 [57.0, 90.3] | 71.2 [55.0, 90.3] |
| **Education, years** | | | | | | |
| Mean (SD) | 15.8 (2.36) | 16.6 (2.48) | 16.3 (2.67) | 16.2 (2.50) | 16.6 (2.36) | 16.4 (2.49) |
| Median [Min, Max] | 16.0 [12.0, 20.0] | 17.0 [8.00, 20.0] | 16.0 [12.0, 20.0] | 16.0 [8.00, 20.0] | 16.0 [12.0, 20.0] | 16.0 [8.00, 20.0] |
| **Ethnicity** | | | | | | |
| Hisp/Latino | 2 (3.6%) | 11 (5.4%) | 7 (5.1%) | 4 (3.4%) | 7 (3.5%) | 31 (4.4%) |
| Not Hisp/Latino | 53 (96.4%) | 191 (94.6%) | 131 (94.9%) | 113 (96.6%) | 191 (96.5%) | 679 (95.6%) |
| **Race/Ethnicity** | | | | | | |
| Asian | 1 (1.8%) | 3 (1.5%) | 1 (0.7%) | 1 (0.9%) | 6 (3.0%) | 12 (1.7%) |
| Black | 2 (3.6%) | 11 (5.4%) | 1 (0.7%) | 6 (5.1%) | 14 (7.1%) | 34 (4.8%) |
| Hispanic/Latino | 2 (3.6%) | 11 (5.4%) | 7 (5.1%) | 4 (3.4%) | 7 (3.5%) | 31 (4.4%) |
| White | 50 (90.9%) | 175 (86.6%) | 126 (91.3%) | 105 (89.7%) | 170 (85.9%) | 626 (88.2%) |
| Multiracial/Other Non-Hispanic | 0 (0%) | 2 (1.0%) | 3 (2.2%) | 1 (0.9%) | 1 (0.5%) | 7 (1.0%) |
| **Marital status** | | | | | | |
| Divorced | 0 (0%) | 26 (12.9%) | 10 (7.2%) | 14 (12.0%) | 26 (13.1%) | 76 (10.7%) |
| Married | 47 (85.5%) | 143 (70.8%) | 117 (84.8%) | 89 (76.1%) | 143 (72.2%) | 539 (75.9%) |
| Never married | 4 (7.3%) | 11 (5.4%) | 3 (2.2%) | 1 (0.9%) | 8 (4.0%) | 27 (3.8%) |
| Widowed | 4 (7.3%) | 22 (10.9%) | 8 (5.8%) | 13 (11.1%) | 21 (10.6%) | 68 (9.6%) |

Values are n (%) unless otherwise specified. CC, complete case; AD, Alzheimer's disease; CN, cognitively normal; EMCI, early mild cognitive impairment; LMCI, late mild cognitive impairment; SMC, significant memory concern; SD, standard deviation.

**Table S2:** Comparison of covariate-adjusted approximate intraclass correlation coefficients (ICCs) between CC and imputed analyses

| **Braak** | **ICC (CC)** | **ICC (Imp)** |
|---|---|---|
| *Method: Raw* | | |
| 1 | 0.135 | 0.106 |
| 2 | 0.169 | 0.143 |
| 3 | 0.101 | 0.095 |
| 4 | 0.109 | 0.096 |
| 5 | 0.114 | 0.121 |
| 6 | 0.161 | 0.177 |
| *Method: ComBat* | | |
| 1 | 0.028 | 0.022 |
| 2 | 0.024 | 0.025 |
| 3 | 0.016 | 0.013 |
| 4 | 0.018 | 0.013 |
| 5 | 0.021 | 0.018 |
| 6 | 0.033 | 0.025 |
| *Method: ComBat Null* | | |
| 1 | 0.051 | 0.033 |
| 2 | 0.042 | 0.036 |
| 3 | 0.030 | 0.022 |
| 4 | 0.018 | 0.015 |
| 5 | 0.024 | 0.022 |
| 6 | 0.035 | 0.034 |
| *Method: QN* | | |
| 1 | 0.037 | 0.032 |
| 2 | 0.021 | 0.016 |
| 3 | 0.043 | 0.043 |
| 4 | 0.032 | 0.033 |
| 5 | 0.036 | 0.041 |
| 6 | 0.019 | 0.023 |
| *Method: QR* | | |
| 1 | 0.060 | 0.048 |
| 2 | 0.054 | 0.041 |
| 3 | 0.050 | 0.050 |
| 4 | 0.044 | 0.033 |
| 5 | 0.069 | 0.037 |
| 6 | 0.043 | 0.029 |

Approximate ICCs were calculated from covariate-adjusted linear model residuals for CC ($n = 710$) and imputed ($n = 986$) analyses, as defined in the Methods. Lower values indicate less dispersion of site mean residuals relative to within-site residual variability. For imputed analyses, estimates were calculated separately within each of the five imputed datasets and then averaged. CC, complete case; Imp, imputed; DM, cognitively impaired group; CU, cognitively unimpaired group; QN, quantile normalization; QR, quantile regression.

**Table S3:** Comparison of effect sizes (Hedges' g) between CC and imputed analyses

| Braak | g (CC) | g (Imp) | Diff | % Change | Sig | SE Ratio | FMI |
|---|---|---|---|---|---|---|---|
| *Method: Raw* | | | | | | | |
| 1 | 0.834 | 0.848 | 0.014 | 1.671 | ✓ | 0.887 | 0.085 |
| 2 | 0.506 | 0.501 | −0.005 | −0.895 | ✓ | 0.909 | 0.132 |
| 3 | 0.709 | 0.712 | 0.003 | 0.369 | ✓ | 0.871 | 0.053 |
| 4 | 0.643 | 0.643 | 0.000 | −0.054 | ✓ | 0.872 | 0.057 |
| 5 | 0.537 | 0.548 | 0.011 | 2.118 | ✓ | 0.878 | 0.067 |
| 6 | 0.325 | 0.339 | 0.014 | 4.294 | ✓ | 0.895 | 0.103 |
| *Method: ComBat* | | | | | | | |
| 1 | 0.832 | 0.838 | 0.007 | 0.809 | ✓ | 0.890 | 0.092 |
| 2 | 0.516 | 0.509 | −0.007 | −1.426 | ✓ | 0.899 | 0.113 |
| 3 | 0.693 | 0.697 | 0.004 | 0.626 | ✓ | 0.873 | 0.057 |
| 4 | 0.611 | 0.631 | 0.021 | 3.369 | ✓ | 0.892 | 0.095 |
| 5 | 0.512 | 0.549 | 0.037 | 7.177 | ✓ | 0.895 | 0.100 |
| 6 | 0.305 | 0.341 | 0.036 | 11.972 | ✓ | 0.897 | 0.106 |
| *Method: ComBat Null* | | | | | | | |
| 1 | 0.755 | 0.793 | 0.038 | 5.043 | ✓ | 0.896 | 0.099 |
| 2 | 0.426 | 0.444 | 0.018 | 4.270 | ✓ | 0.881 | 0.074 |
| 3 | 0.656 | 0.687 | 0.031 | 4.774 | ✓ | 0.869 | 0.044 |
| 4 | 0.556 | 0.596 | 0.041 | 7.310 | ✓ | 0.878 | 0.063 |
| 5 | 0.466 | 0.530 | 0.064 | 13.676 | ✓ | 0.888 | 0.083 |
| 6 | 0.221 | 0.293 | 0.072 | 32.398 | ✓ | 0.895 | 0.099 |
| *Method: QN* | | | | | | | |
| 1 | 0.687 | 0.741 | 0.054 | 7.884 | ✓ | 0.898 | 0.103 |
| 2 | 0.463 | 0.450 | −0.013 | −2.766 | ✓ | 0.885 | 0.086 |
| 3 | 0.641 | 0.679 | 0.038 | 5.935 | ✓ | 0.875 | 0.057 |
| 4 | 0.551 | 0.574 | 0.023 | 4.116 | ✓ | 0.871 | 0.050 |
| 5 | 0.424 | 0.526 | 0.102 | 24.093 | ✓ | 0.893 | 0.090 |
| 6 | 0.252 | 0.326 | 0.074 | 29.591 | ✓ | 0.894 | 0.098 |
| *Method: QR* | | | | | | | |
| 1 | 0.878 | 0.881 | 0.003 | 0.318 | ✓ | 0.894 | 0.100 |
| 2 | 0.543 | 0.521 | −0.023 | −4.163 | ✓ | 0.892 | 0.100 |
| 3 | 0.737 | 0.719 | −0.018 | −2.466 | ✓ | 0.861 | 0.035 |
| 4 | 0.655 | 0.643 | −0.012 | −1.828 | ✓ | 0.889 | 0.093 |
| 5 | 0.587 | 0.558 | −0.029 | −4.961 | ✓ | 0.910 | 0.136 |
| 6 | 0.350 | 0.340 | −0.010 | −2.837 | ✓ | 0.906 | 0.127 |

Effect sizes (Hedges' g) for DM vs CU group discrimination comparing CC ($n = 710$) and imputed ($n = 986$) analyses. Diff = g(Imp) - g(CC); % Change = (Diff / g(CC)) × 100. $SE\,Ratio = SE_{Imp}/SE_{CC}$, where values smaller than 1 indicate improved precision with imputation. ✓ indicates statistical significance ($p < 0.001$) maintained in both analyses. All effect sizes were calculated using Hedges' correction for small sample bias. Estimates from five multiply imputed datasets were pooled using Rubin's rules. CC, complete case; Imp, imputed; DM, cognitively impaired group; CU, cognitively unimpaired group; QN, quantile normalization; QR, quantile regression; SE, standard error; FMI, fraction of missing information.

**Table S4:** Comparison of covariate-adjusted regression coefficients ($\hat{\beta}_1$; DM vs CU) between CC and imputed analyses across harmonization methods and Braak regions

| Braak | $\hat{\beta}_1$ (CC) | $\hat{\beta}_1$ (Imp) | Diff | % Change | SE Ratio | FMI |
|---|---|---|---|---|---|---|
| *Method: Raw* | | | | | | |
| 1 | 0.1779 | 0.1797 | 0.0018 | 1.018 | 0.942 | 0.1628 |
| 2 | 0.0809 | 0.0780 | −0.0028 | −3.494 | 0.947 | 0.1629 |
| 3 | 0.1371 | 0.1362 | −0.0009 | −0.693 | 0.891 | 0.0605 |
| 4 | 0.1121 | 0.1079 | −0.0042 | −3.763 | 0.891 | 0.0966 |
| 5 | 0.0958 | 0.0918 | −0.0040 | −4.151 | 0.858 | 0.0926 |
| 6 | 0.0418 | 0.0412 | −0.0006 | −1.473 | 0.892 | 0.1223 |
| *Method: ComBat* | | | | | | |
| 1 | 0.1666 | 0.1696 | 0.0029 | 1.763 | 0.988 | 0.2161 |
| 2 | 0.0760 | 0.0738 | −0.0022 | −2.893 | 0.989 | 0.2168 |
| 3 | 0.1191 | 0.1202 | 0.0012 | 0.970 | 0.944 | 0.1458 |
| 4 | 0.0959 | 0.0975 | 0.0016 | 1.693 | 0.948 | 0.1778 |
| 5 | 0.0805 | 0.0826 | 0.0020 | 2.537 | 0.906 | 0.1598 |
| 6 | 0.0348 | 0.0375 | 0.0027 | 7.684 | 0.925 | 0.1555 |
| *Method: ComBat Null* | | | | | | |
| 1 | 0.1503 | 0.1605 | 0.0102 | 6.779 | 0.975 | 0.2126 |
| 2 | 0.0616 | 0.0637 | 0.0020 | 3.314 | 0.949 | 0.1732 |
| 3 | 0.1110 | 0.1177 | 0.0068 | 6.096 | 0.934 | 0.1329 |
| 4 | 0.0862 | 0.0919 | 0.0057 | 6.555 | 0.928 | 0.1494 |
| 5 | 0.0740 | 0.0815 | 0.0074 | 10.044 | 0.899 | 0.1464 |
| 6 | 0.0270 | 0.0345 | 0.0075 | 27.863 | 0.926 | 0.1606 |
| *Method: QN* | | | | | | |
| 1 | 0.1161 | 0.1270 | 0.0108 | 9.313 | 0.966 | 0.1966 |
| 2 | 0.0562 | 0.0540 | −0.0022 | −3.941 | 0.970 | 0.1858 |
| 3 | 0.0919 | 0.0992 | 0.0073 | 7.972 | 0.933 | 0.1430 |
| 4 | 0.0721 | 0.0739 | 0.0018 | 2.485 | 0.915 | 0.1147 |
| 5 | 0.0565 | 0.0707 | 0.0142 | 25.160 | 0.934 | 0.1851 |
| 6 | 0.0242 | 0.0325 | 0.0083 | 34.182 | 0.935 | 0.1551 |
| *Method: QR* | | | | | | |
| 1 | 0.1182 | 0.1125 | −0.0058 | −4.890 | 0.990 | 0.3132 |
| 2 | 0.0610 | 0.0542 | −0.0067 | −11.013 | 0.909 | 0.1529 |
| 3 | 0.0966 | 0.0756 | −0.0211 | −21.791 | 0.774 | 0.1402 |
| 4 | 0.0808 | 0.0675 | −0.0133 | −16.504 | 0.786 | 0.0602 |
| 5 | 0.0712 | 0.0616 | −0.0095 | −13.385 | 0.873 | 0.1495 |
| 6 | 0.0355 | 0.0326 | −0.0029 | −8.188 | 0.872 | 0.1008 |

Regression coefficients ($\hat{\beta}_1$) represent covariate-adjusted mean tau SUVR differences between DM and CU groups. Diff $= \hat{\beta}_{1,\text{Imp}} - \hat{\beta}_{1,\text{CC}}$; % Change $= (\text{Diff}/\hat{\beta}_{1,\text{CC}}) \times 100$. SE Ratio $= SE_{\text{Imp}}/SE_{\text{CC}}$. Estimates from five multiply imputed datasets were pooled using Rubin's rules. CC, complete case; Imp, imputed; DM, cognitively impaired group; CU, cognitively unimpaired group; QN, quantile normalization; QR, quantile regression; SE, standard error; FMI, fraction of missing information.

**Table S5:** Comparison of statistical significance for disease group effects between CC and imputed analyses

| Braak | $p$ (CC) | $p$ (Imp) | $-\log_{10}(p_{CC})$ | $-\log_{10}(p_{Imp})$ | $\Delta - \log_{10}(p)$ |
|---|---|---|---|---|---|
| *Method: Raw* | | | | | |
| 1 | $2.71{\times}10^{-20}$ | $6.47{\times}10^{-19}$ | 19.6 | 18.2 | $-1.38$ |
| 2 | $2.39{\times}10^{-8}$ | $4.79{\times}10^{-8}$ | 7.6 | 7.3 | $-0.30$ |
| 3 | $2.26{\times}10^{-14}$ | $1.25{\times}10^{-17}$ | 13.6 | 16.9 | 3.26 |
| 4 | $1.63{\times}10^{-12}$ | $5.79{\times}10^{-14}$ | 11.8 | 13.2 | 1.45 |
| 5 | $2.85{\times}10^{-9}$ | $5.13{\times}10^{-11}$ | 8.5 | 10.3 | 1.75 |
| 6 | $5.50{\times}10^{-5}$ | $1.09{\times}10^{-5}$ | 4.3 | 5.0 | 0.70 |
| *Method: ComBat* | | | | | |
| 1 | $4.98{\times}10^{-20}$ | $1.79{\times}10^{-15}$ | 19.3 | 14.7 | $-4.56$ |
| 2 | $1.35{\times}10^{-8}$ | $2.19{\times}10^{-7}$ | 7.9 | 6.7 | $-1.21$ |
| 3 | $2.11{\times}10^{-13}$ | $1.18{\times}10^{-13}$ | 12.7 | 12.9 | 0.25 |
| 4 | $3.55{\times}10^{-11}$ | $4.46{\times}10^{-11}$ | 10.4 | 10.4 | $-0.10$ |
| 5 | $1.12{\times}10^{-8}$ | $8.33{\times}10^{-10}$ | 8.0 | 9.1 | 1.13 |
| 6 | $9.94{\times}10^{-5}$ | $1.00{\times}10^{-5}$ | 4.0 | 5.0 | 1.00 |
| *Method: ComBat Null* | | | | | |
| 1 | $3.86{\times}10^{-16}$ | $2.17{\times}10^{-14}$ | 15.4 | 13.7 | $-1.75$ |
| 2 | $6.69{\times}10^{-6}$ | $2.30{\times}10^{-6}$ | 5.2 | 5.6 | 0.46 |
| 3 | $7.05{\times}10^{-12}$ | $1.04{\times}10^{-13}$ | 11.2 | 13.0 | 1.83 |
| 4 | $3.08{\times}10^{-9}$ | $9.11{\times}10^{-11}$ | 8.5 | 10.0 | 1.53 |
| 5 | $1.51{\times}10^{-7}$ | $7.57{\times}10^{-10}$ | 6.8 | 9.1 | 2.30 |
| 6 | $2.78{\times}10^{-3}$ | $5.63{\times}10^{-5}$ | 2.6 | 4.2 | 1.69 |
| *Method: QN* | | | | | |
| 1 | $3.34{\times}10^{-13}$ | $2.61{\times}10^{-13}$ | 12.5 | 12.6 | 0.11 |
| 2 | $1.64{\times}10^{-6}$ | $5.04{\times}10^{-6}$ | 5.8 | 5.3 | $-0.49$ |
| 3 | $4.76{\times}10^{-11}$ | $5.50{\times}10^{-13}$ | 10.3 | 12.3 | 1.94 |
| 4 | $5.79{\times}10^{-9}$ | $1.82{\times}10^{-10}$ | 8.2 | 9.7 | 1.50 |
| 5 | $2.10{\times}10^{-6}$ | $3.28{\times}10^{-9}$ | 5.7 | 8.5 | 2.81 |
| 6 | $1.28{\times}10^{-3}$ | $6.97{\times}10^{-6}$ | 2.9 | 5.2 | 2.26 |
| *Method: QR* | | | | | |
| 1 | $3.56{\times}10^{-22}$ | $5.95{\times}10^{-12}$ | 21.4 | 11.2 | $-10.22$ |
| 2 | $1.72{\times}10^{-9}$ | $1.34{\times}10^{-8}$ | 8.8 | 7.9 | $-0.89$ |
| 3 | $3.57{\times}10^{-16}$ | $6.11{\times}10^{-15}$ | 15.4 | 14.2 | $-1.23$ |
| 4 | $2.73{\times}10^{-13}$ | $6.01{\times}10^{-15}$ | 12.6 | 14.2 | 1.66 |
| 5 | $3.74{\times}10^{-11}$ | $3.14{\times}10^{-10}$ | 10.4 | 9.5 | $-0.92$ |
| 6 | $9.42{\times}10^{-6}$ | $3.58{\times}10^{-6}$ | 5.0 | 5.4 | 0.42 |

Statistical significance for DM vs CU group comparisons from CC ($n = 710$) and imputed ($n = 986$) analyses. $\Delta - \log_{10}(p) = -\log_{10}(p_{MI}) - -\log_{10}(p_{CC})$ represents the change in statistical strength after imputation. Positive values indicate stronger statistical evidence in the imputed analyses. All comparisons remained statistically significant ($p < 0.05$). Estimates from five multiply imputed datasets were pooled using Rubin's rules. CC, complete case; Imp, imputed; DM, cognitively impaired group; CU, cognitively unimpaired group; QN, quantile normalization; QR, quantile regression.

### S2 Supplementary Figures

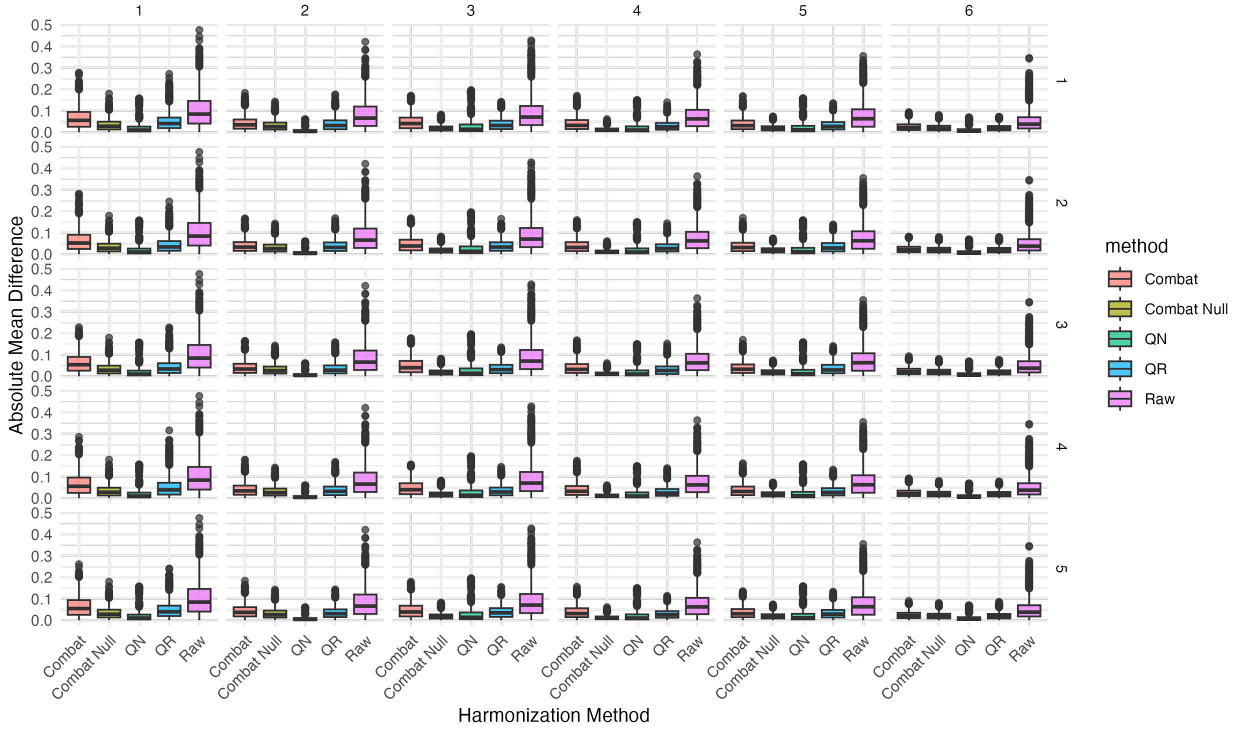


**Figure S1:** Distribution of pairwise site mean differences in tau SUVR across five multiply imputed datasets. Boxplots display absolute differences in site-specific mean SUVR for each harmonization method and Braak region. Pairwise differences were computed separately within each imputed dataset and are shown for each of the five imputations to evaluate the stability of site-level harmonization across imputations. Lower values and narrower distributions indicate reduced residual site heterogeneity. QN, quantile normalization; QR, quantile regression; SUVR, standardized uptake value ratio.

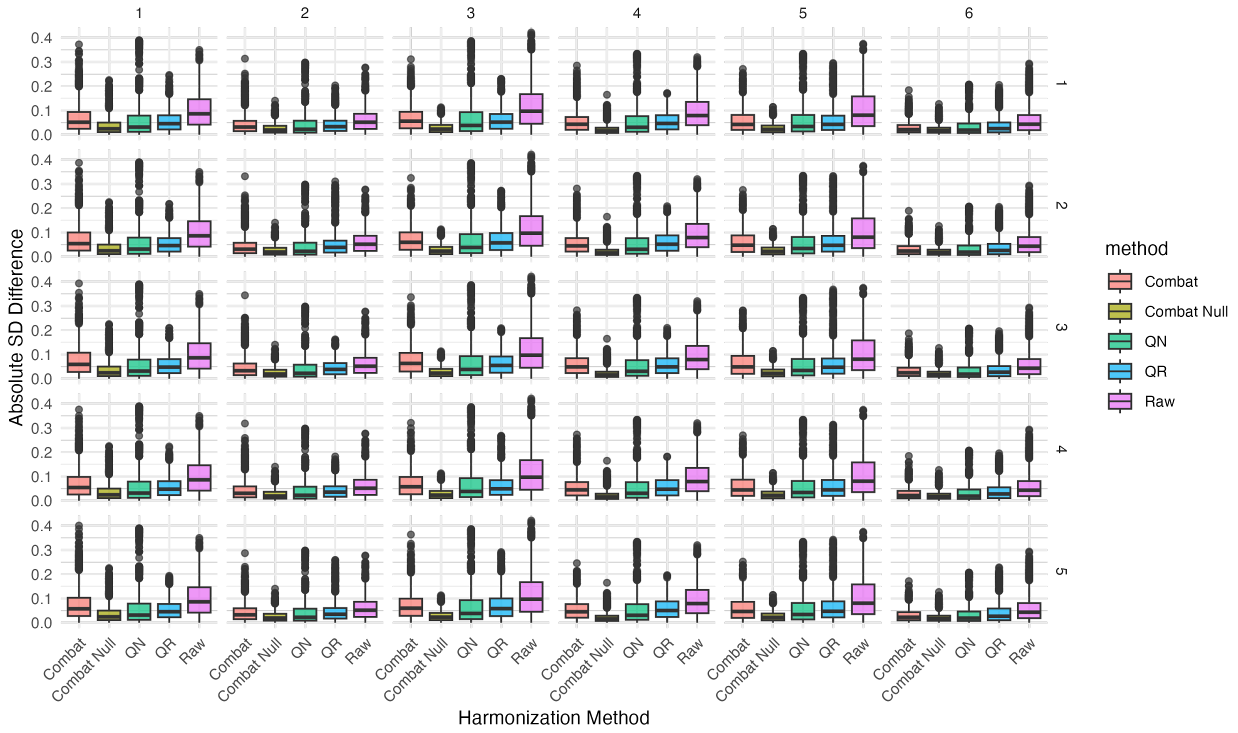


**Figure S2:** Distribution of pairwise site differences in within-site standard deviations of tau SUVR across five multiply imputed datasets. Boxplots display absolute differences in site-specific SUVR variability between all site pairs for each harmonization method and Braak region. Pairwise SD differences were computed separately within each imputed dataset and are shown for each of the five imputations to evaluate the stability of variance harmonization across imputations. Lower values indicate reduced heterogeneity in within-site SUVR variability across scanning locations. QN, quantile normalization; QR, quantile regression; SUVR, standardized uptake value ratio.

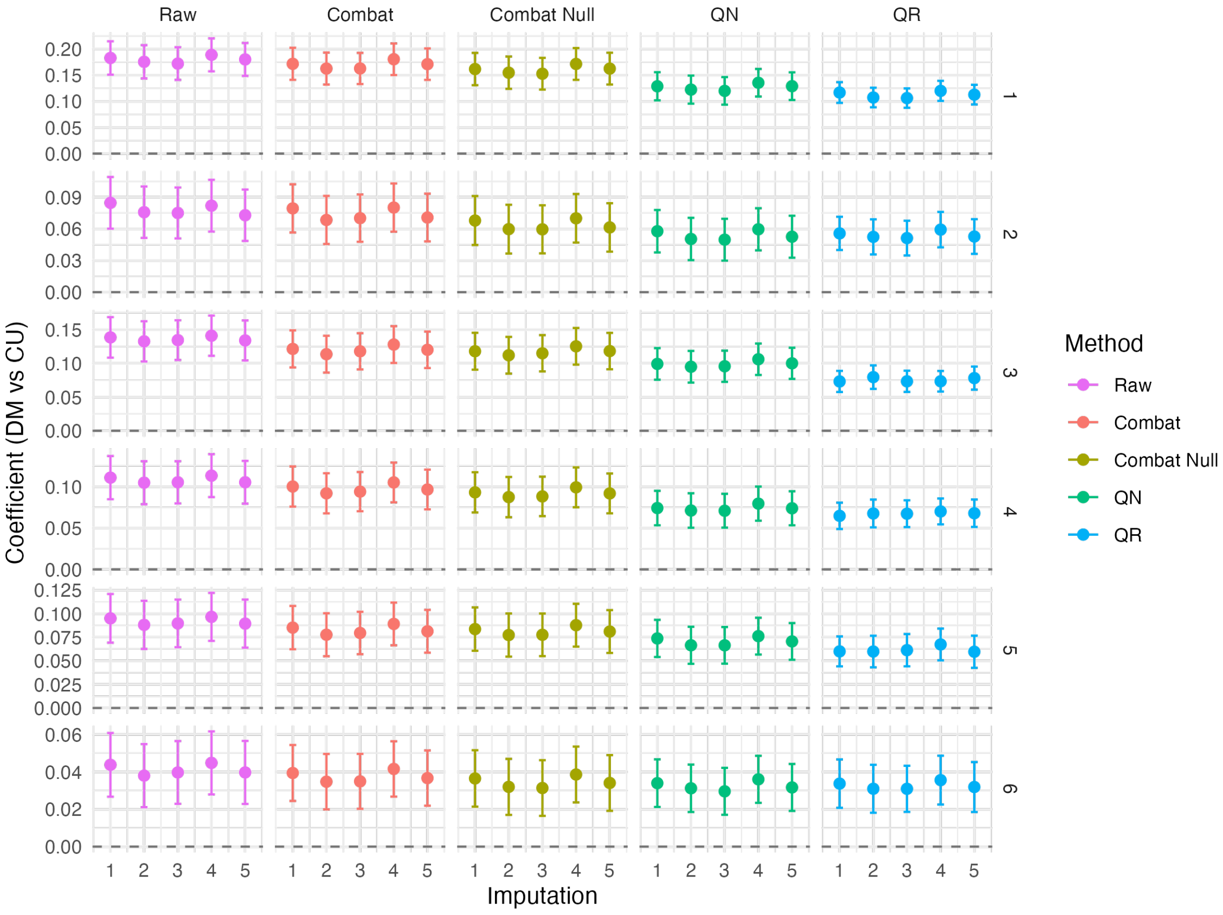


**Figure S3:** Group effect estimates for diagnosis across five multiply imputed datasets. Points represent estimates from individual imputations with 95% confidence intervals. DM, cognitively impaired group; CU, cognitively unimpaired group; QN, quantile normalization; QR, quantile regression.

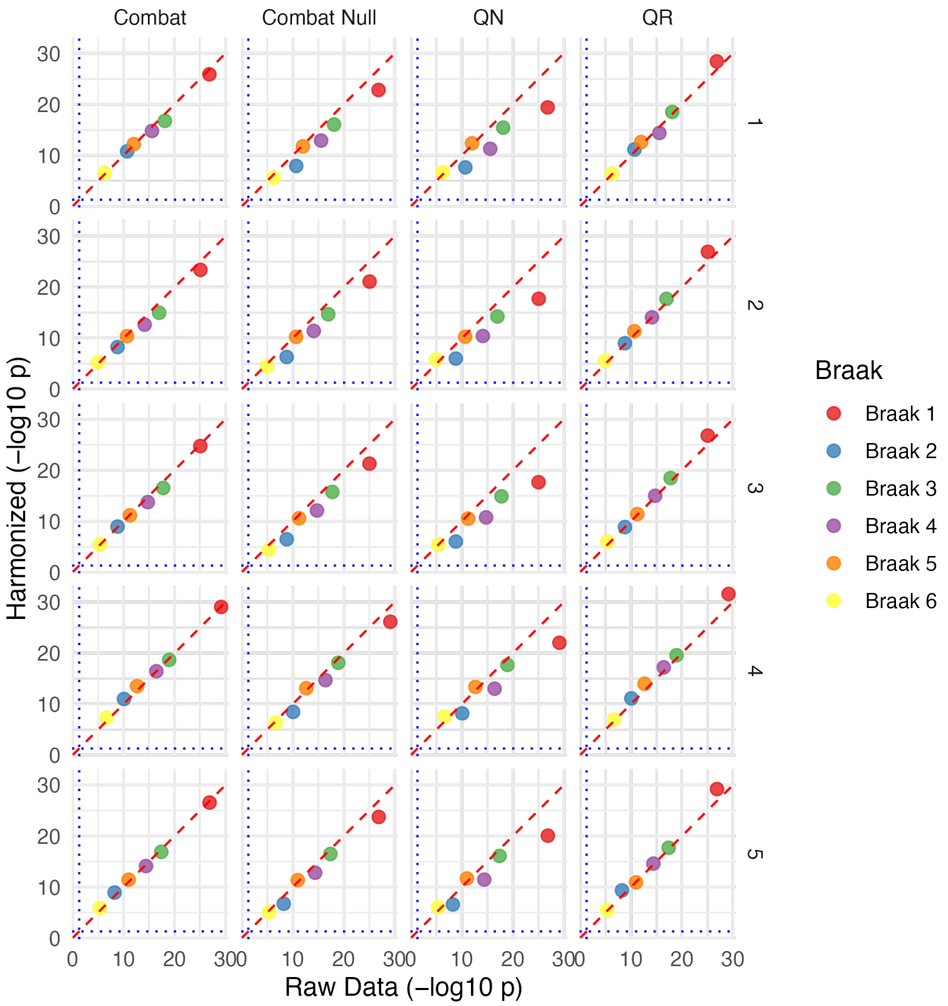


**Figure S4:** Comparison of statistical significance between raw and harmonized data across methods and Braak regions for five imputed datasets. Scatter plots display $-\log_{10}(p)$ values for DM vs CU group effects in raw data (x-axis) versus harmonized data (y-axis) for ComBat, ComBat Null, QN, and QR. The red dashed diagonal indicates perfect agreement, while blue dotted lines denote the significance threshold ($p = 0.05$) on both axes. Points above both thresholds indicate statistical significance in both analyses. All group comparisons remained statistically significant ($p < 0.001$) in both raw and harmonized data. DM, cognitively impaired group; CU, cognitively unimpaired group; QN, quantile normalization; QR, quantile regression.

### S3 Sensitivity analysis using meta-temporal SUVR

**Methods.** We evaluated meta-temporal standardized uptake value ratio (SUVR) using the existing `META_TEMPORAL_SUVR` variable, retaining the original eligibility criteria, site filters, and complete-case (CC) cohort. Missing covariates were multiply imputed using the same settings as the main analysis (five imputations, 50 iterations), with meta-temporal SUVR included as an observed predictor. Imaging outcomes were not imputed.

ComBat and ComBat Null were fitted jointly to the six Braak composites and meta-temporal SUVR, allowing empirical-Bayes borrowing across seven features. Quantile normalization (QN) and quantile regression (QR) were applied separately to the meta-temporal feature. Harmonization was performed separately for the CC dataset and each imputed dataset. Only meta-temporal results are reported here; the main-paper analyses were unchanged.

We evaluated absolute pairwise site mean and standard deviation (SD) differences, adjusted approximate intraclass correlation coefficients (ICCs), Hedges' $g$, and adjusted regression coefficients comparing the cognitively impaired (DM) and cognitively unimpaired (CU) groups. Regression adjusted for age, sex, education, APOE4, race/ethnicity, and marital status. Effect estimates from the imputed analyses (Imp) were pooled using Rubin's rules with Barnard–Rubin-adjusted degrees of freedom. Precision was evaluated using standard error (SE) ratios and the fraction of missing information (FMI).

**Results.** The CC analysis included 710 participants from 53 sites, and the imputed analyses included 986 participants from 57 sites (Table S6). All harmonization methods reduced median absolute site mean and SD differences and adjusted approximate ICCs relative to raw data within each analysis. Positive DM–CU differences remained under every method (all $p < 0.001$). Hedges' $g$ ranged from 0.612 to 0.731 in CC and from 0.676 to 0.739 in the imputed analyses. Pooled SEs were smaller than CC SEs, with Imp-to-CC ratios of 0.883–0.904 for Hedges' $g$ and 0.760–0.936 for regression coefficients. QR showed the largest CC–Imp coefficient change (0.113 versus 0.087). These results indicate that, for meta-temporal SUVR, harmonization reduced site variability while retaining biological group differences, and imputation yielded smaller estimated SEs. Effect magnitudes varied by method, and QR estimates require caution because the fitting algorithm reported potentially nonunique solutions.

**Table S6:** Site variability and biological group differences in meta-temporal SUVR: comparison of CC and imputed analyses.

**A. Site variability and biological group differences**

| Dataset | Method | Site mean diff. | Site SD diff. | Adj. ICC | Hedges' $g$ [95% CI] | $\hat{\beta}_1$ [95% CI] |
|---|---|---|---|---|---|---|
| CC | Raw | 0.094 | 0.130 | 0.105 | 0.731 [0.578, 0.885] | 0.181 [0.138, 0.225] |
| CC | ComBat | 0.058 | 0.089 | 0.025 | 0.698 [0.546, 0.851] | 0.155 [0.115, 0.194] |
| CC | ComBat Null | 0.027 | 0.031 | 0.031 | 0.661 [0.509, 0.814] | 0.144 [0.105, 0.184] |
| CC | QN | 0.018 | 0.048 | 0.043 | 0.612 [0.460, 0.763] | 0.114 [0.080, 0.148] |
| CC | QR | 0.043 | 0.079 | 0.049 | 0.723 [0.570, 0.876] | 0.113 [0.086, 0.140] |
| Imp | Raw | 0.089 | 0.123 | 0.092 | 0.739 [0.602, 0.877] | 0.178 [0.137, 0.218] |
| Imp | ComBat | 0.054 | 0.076 | 0.021 | 0.716 [0.577, 0.855] | 0.157 [0.119, 0.194] |
| Imp | ComBat Null | 0.025 | 0.026 | 0.022 | 0.703 [0.568, 0.838] | 0.153 [0.117, 0.188] |
| Imp | QN | 0.017 | 0.044 | 0.041 | 0.676 [0.541, 0.811] | 0.125 [0.095, 0.156] |
| Imp | QR | 0.035 | 0.060 | 0.049 | 0.739 [0.603, 0.874] | 0.087 [0.066, 0.108] |

**B. Precision and missing information**

| Method | SE ratio ($g$) | SE ratio ($\hat{\beta}_1$) | FMI ($g$) | FMI ($\hat{\beta}_1$) |
|---|---|---|---|---|
| Raw | 0.895 | 0.928 | 0.111 | 0.169 |
| ComBat | 0.904 | 0.936 | 0.129 | 0.159 |
| ComBat Null | 0.884 | 0.907 | 0.081 | 0.103 |
| QN | 0.887 | 0.895 | 0.084 | 0.083 |
| QR | 0.883 | 0.760 | 0.082 | 0.255 |

Site differences are medians of absolute pairwise differences. For imputed analyses, absolute differences were first averaged across imputations within each site pair. CC and imputed analyses included different site sets. Adjusted approximate ICCs were calculated from covariate-adjusted residuals and averaged across imputations; they are reported descriptively. Hedges' $g$ and $\hat{\beta}_1$ represent DM–CU contrasts. All two-sided $p$-values for these contrasts were $< 0.001$, without adjustment for multiplicity. CC confidence intervals for $g$ use a normal approximation; CC regression intervals and pooled imputed intervals use $t$ inference. SE ratio $= SE_{\text{Imp}}/SE_{\text{CC}}$. FMI includes the finite-sample adjustment. The implemented imputation model excluded site as a predictor. QR fitting produced warnings of potentially nonunique solutions.

CC, complete case; Imp, imputed; DM, cognitively impaired group; CU, cognitively unimpaired group; SUVR, standardized uptake value ratio; SD, standard deviation; ICC, intraclass correlation coefficient; CI, confidence interval; QN, quantile normalization; QR, quantile regression; SE, standard error; FMI, fraction of missing information.
